# A comprehensive benchmark of transcriptome-wide fusion detection using long-read RNA sequencing

**DOI:** 10.64898/2026.08.07.743439

**Authors:** Ryley Dorney, Siyuan Wu, Julia Yun-Hsuan Hung, Lionel Hebbard, Ulf Schmitz

**Affiliations:** Computational BioMedicine Lab, College of Science and Engineering, James Cook University, Douglas, QLD 4811, Australia; Centre for Tropical Bioinformatics and Molecular Biology, Australian Institute of Tropical Health and Medicine, James Cook University, Cairns 4878, Australia; Department of Biomedical Sciences and Molecular and Cell Biology, College of Medicine and Dentistry, College of Science and Engineering, James Cook University, Townsville, QLD, Australia; Storr Liver Centre, Westmead Institute for Medical Research, Westmead Hospital and University of Sydney, Sydney, NSW, Australia; Australian Institute for Tropical Health and Medicine, Townsville, QLD 4814, Australia; Centenary Institute, The University of Sydney, Camperdown, NSW 2050, Australia

## Abstract

Fusion transcripts contribute to cancer, inherited diseases, developmental disorders, and evolution. Long-read RNA sequencing enables direct sequencing of full-length transcripts, creating new opportunities to detect complex fusion architectures, including previously inaccessible multi-segmented fusion transcripts. However, accurate transcriptome-wide fusion detection remains challenging because existing methods struggle to distinguish genuine fusion events from technical artefacts.

Here, we present a comprehensive benchmark of transcriptome-wide fusion detection using simulated datasets and transcriptomes from three cancer cell lines across Oxford Nanopore Technologies (ONT) cDNA, PCR-cDNA, and direct RNA sequencing, Pacific Biosciences (PacBio) Kinnex sequencing, Illumina short-read RNA sequencing, six long-read fusion callers, and multiple analysis strategies.

False-positive fusion calls remained the dominant limitation across sequencing platforms and algorithms. Increasing sequencing depth improved recall but also amplified spurious fusion calls, whereas higher read-support thresholds improved precision at the expense of sensitivity. ONT PCR-cDNA sequencing combined with *CTAT-LR*-*Fusion* achieved the best overall balance between precision and recall, whereas *JAFFAL* was the only caller to reliably identify simulated tri-gene fusions. Consensus calling reduced false positives but markedly reduced sensitivity, with only one of 400 simulated fusions detected by all six callers. Breakpoint localisation emerged as a major limitation across all methods.

Long-read sequencing consistently recovered more validated fusion transcripts than short-read sequencing, enabled detection of complex tri-gene fusions, and produced more biologically plausible fusion landscapes with fewer promiscuous gene partners. Collectively, our results establish the first comprehensive benchmarking framework for transcriptome-wide fusion detection, using long-read RNA sequencing, and provide practical guidance for selecting sequencing workflows and computational strategies, while identifying key priorities for future algorithm development.

## Introduction

Fusion transcripts are chimeric RNA molecules comprising exons, and sometimes introns, derived from two or more parental genes (1). They commonly arise from genomic rearrangements that generate fusion genes, although they can also result from transcriptional mechanisms such as *trans*-splicing or read-through transcription. Fusion transcripts are important molecular drivers of human diseases, particularly cancer, where they are estimated to contribute to 16.5% of malignancies and serve as the sole driver in more than 1% of cases (2). Beyond cancer, germline gene fusion events have been associated with a range of inherited disorders, including red-green colour blindness (3), dilated cardiomyopathy (4), cerebral malformation (5), and various rare genetic diseases (6, 7). Fusion genes and transcripts have also contributed to evolutionary innovation across diverse taxa, including plants (8), fungi (9), and primates (10).

Fusion transcripts are not exclusively generated by genomic rearrangements. Alternative mechanisms include *cis*- and *trans*-splicing between neighbouring or distant genes, read-through transcription, sense-antisense fusions, and other complex transcriptional events (1). Even among rearrangement-derived fusions, transcript architecture can be considerably more intricate than simple two-gene chimeras. Several studies have described *bridged* and *multi-hop* fusions, in which multiple structural rearrangements juxtapose more than two genomic loci (9, 11–16). Although the intervening bridged sequence is typically removed during RNA splicing, mature transcripts comprising exons from three distinct genes have been reported, demonstrating that genuine tri-gene fusion transcripts can arise in human cells (17, 18). The growing recognition of these complex fusion architectures presents new challenges for computational fusion detection and highlights the need for sequencing technologies capable of resolving full-length transcript structures.

Transcriptome-wide fusion discovery depends on four interacting components: sequencing technology, library preparation, computational algorithms, and analysis strategies. However, short-read sequencing remains inherently limited in resolving complex transcript architectures (19). Individual reads rarely span multiple fusion junctions, preventing comprehensive reconstruction of multi-segmented fusion transcripts, while most short-read fusion callers identify breakpoints independently rather than linking them into complete transcript structures. Consequently, the prevalence, biological significance, and diversity of complex fusion transcripts remain poorly understood.

Long-read sequencing technologies, including those developed by Pacific Biosciences (PacBio) and Oxford Nanopore (ONT), overcome many of these limitations by generating reads that frequently span entire RNA molecules (19). Full-length transcript sequencing enables direct identification of all fusion breakpoints, transcription start and end sites, splice isoforms, and retained exonic or intronic sequences within individual fusion transcripts. Improvements in sequencing accuracy and throughput have accelerated the adoption of long-read RNA sequencing for fusion discovery, creating new opportunities to identify complex and multi-segmented fusion transcripts. However, the performance of current long-read fusion detection algorithms across sequencing platforms and library preparation methods has not been comprehensively evaluated (1).

Recent improvements in sequencing accuracy and throughput have enabled long-read RNA sequencing to detect multi-segment fusions without relying on short-read error correction. Multiple library preparation methods are now available, including PCR-cDNA, direct cDNA and direct RNA sequencing, alongside distinct sequencing platforms from ONT and PacBio.

In parallel, several computational tools have been developed to identify fusion transcripts from long-read RNA-seq data. Despite these advances, their relative performance across sequencing platforms, library preparation methods and experimental conditions remains largely unknown, and no independent benchmark has systematically evaluated their strength and limitations.

Here, we present a comprehensive benchmark of transcriptome-wide fusion detection using long-read RNA sequencing. We systematically evaluated six computational fusion callers across multiple sequencing platforms and library preparation methods, including ONT direct cDNA, PCR-cDNA, and direct RNA sequencing, PacBio Kinnex sequencing, and matched Illumina short-read RNA sequencing. Using both simulated datasets and experimentally derived cancer cell line transcriptomes, we assessed sensitivity, precision, breakpoint localisation, computational performance, sequencing depth, read accuracy, and the detection of complex multi-segmented fusion transcripts. Our study provides practical recommendations for selecting sequencing workflows and computational strategies while defining the current capabilities, limitations, and future priorities for long-read fusion transcript discovery.

## Results

Transcriptome-wide fusion discovery depends on four interacting components: sequencing technology, library preparation, computational algorithms, and downstream analysis strategies. To systematically evaluate the contribution of each component, we established a comprehensive benchmarking framework comprising simulated fusion transcript datasets with known ground truth, publicly available cancer cell line transcriptomes, and newly generated Huh7 long-read and matched Illumina RNA sequencing data **(Figure 1A).** Across these datasets we compared six long-read fusion callers, multiple sequencing platforms and library preparation methods. Specifically, we evaluated (i) the performance of computational fusion callers using simulated datasets, (ii) the influence of sequencing depth, read identity and analysis strategies on fusion detection, (iii) fusion discovery in experimentally derived cancer transcriptomes, and (iv) the impact of long-read library preparation methods on transcriptome-wide fusion discovery **(Figure 1A).**

**Figure 1.**
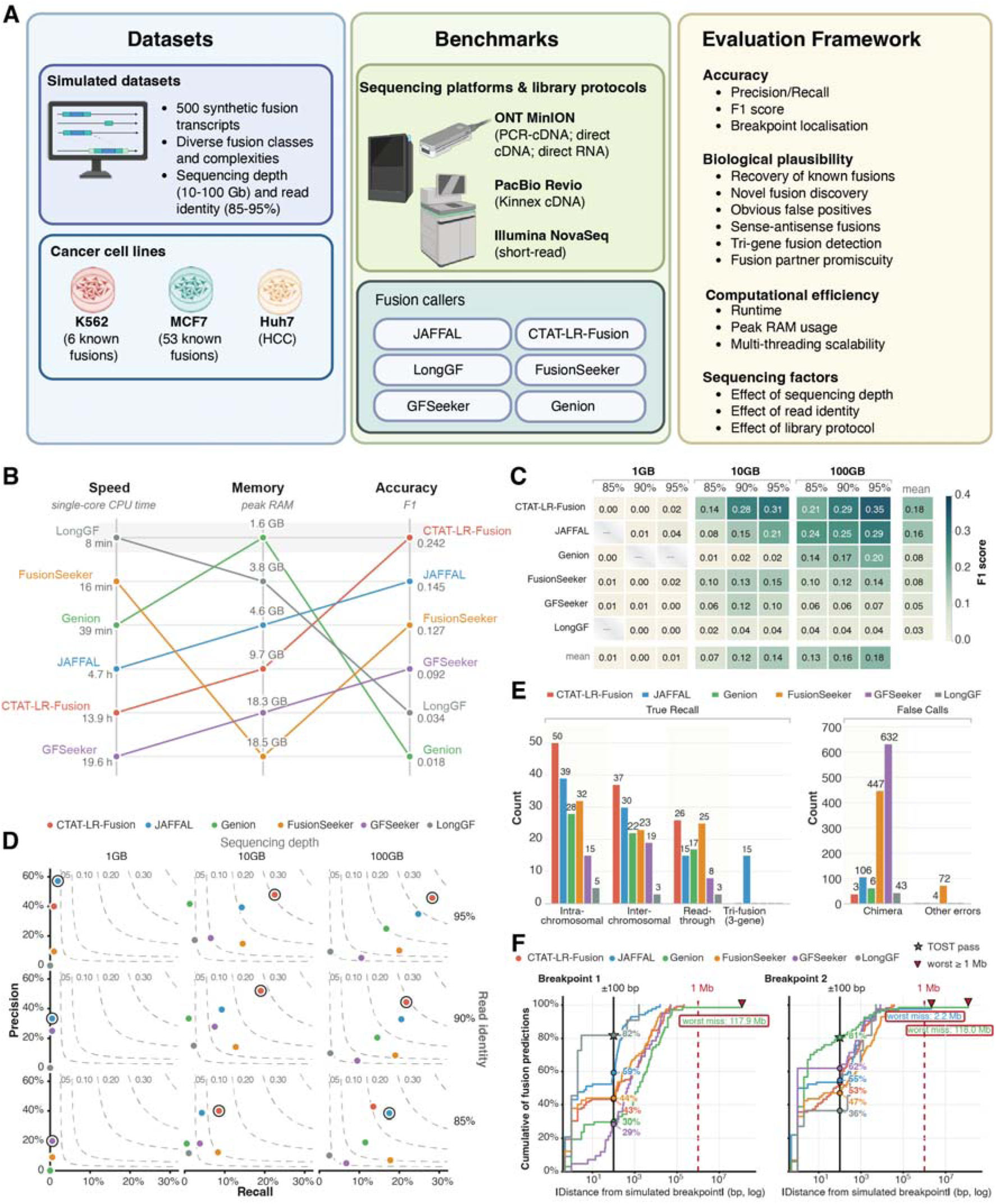
Benchmarking framework. **(A)** We developed a comprehensive benchmarking framework to evaluate transcriptome-wide fusion discovery across sequencing technologies, library preparation methods, computational algorithms, and downstream analysis strategies. **(B)** Computational performance. The six long-read callers are ordered, best at top, on three axes: speed (single-core user CPU time on a 10 Gb simulated dataset), memory (peak RAM), and accuracy (F1 at 10 Gb, averaged over the three sequence-identity levels). Each caller is drawn as a line linking its three ranks. **(C)** F1 scores across sequencing depth and read identify. (D) Precision-recall position across the simulation grid. **(E)** Fusion recall for each fusion subtype (out of 100 simulated fusions each). **(F)** The absolute distance between the breakpoint position predicted by the fusion caller and the simulated breakpoint of spiked-in fusions. 100Gb sequencing depth, 95% mean sequence identity.

### Benchmarking long-read fusion callers using simulated datasets

#### Computational resource usage varies substantially across fusion callers

To systematically evaluate long-read fusion callers, we generated simulated long-read RNA sequencing data containing 500 synthetic fusion transcripts, including 100 tri-gene fusions and 100 read-through transcripts, and with defined sequencing error rates, read lengths, and fusion complexities (**Supplementary Data 1; Supplementary Table 1**). We also tested these tools on long-read RNA sequencing data from MCF7 and K562 cell lines, which contain well-characterised fusion transcripts.

We first compared the computational requirements of each fusion caller using 1GB and 10GB simulated data. Runtime and memory usage varied substantially between tools **(Figure 1B).** *LongGF* consistently exhibited the shortest runtime, followed by *FusionSeeker* on larger datasets, whereas *Genion* was among the fastest on the 1 GB dataset, but recovered comparatively few fusion transcripts (**Supplementary Data 2**). In contrast, *GFSeeker* and *CTAT-LR-Fusion* required substantially longer runtimes than all other methods, despite producing competitive fusion detection performance.

Peak memory usage also differed markedly. *GFSeeker* and *FusionSeeker* consistently required the most memory (16.9-18.5 GB), even when processing the 1 GB datasets, whereas *LongGF*, *Genion* and *JAFFAL* completed analyses using less than 1.2 GB of RAM (**Figure 1B**; **Supplementary Data 2**).

Multi-threading provided limited performance gains for most callers. Increasing the number of CPU cores generally reduced wall time but had little effect on total user time. Notably, *FusionSeeker* and *CTAT-LR*-*Fusion’s* exhibited longer wall times when executed with multiple threads, suggesting inefficient parallelisation. In contrast, *JAFFAL* scaled efficiently with multithreading, approximately halving wall time without increasing memory usage (**Supplementary Data 2**).

#### Fusion callers exhibit limited recall and frequent false-positive predictions

To compare fusion caller performance, we quantified the precision, recall and F1 score of simulated fusion transcripts. Precision was defined as the proportion of correctly identified fusions relative to all reported fusions, excluding false chimeras and partially reconstructed events. Overall performance was lower than expected, with no caller achieving an F1 score above 0.35 (**Figure 1C, Supplementary Figure 1**). Low F1 scores were largely driven by poor recall, which did not exceed 30% for any method, although false-positive fusion calls also reduced precision. Among the evaluated tools, *CTAT-LR-Fusion* achieved the highest overall F1 score, followed by *JAFFAL,* owing largely to their superior precision. *CTAT-LR-Fusion* consistently maintained precision above 40% across all sequencing depths. Moreover, including partially reconstructed fusion transcripts substantially inflated performance estimates **(Supplementary Figure 2),** particularly precision, because a single simulated fusion could generate multiple partial predictions. We therefore excluded partial matches from all subsequent performance analyses.

#### Increasing sequencing depth improves recall but also increases false-positive fusion calls

To assess the effect of sequencing depth on fusion detection, we simulated datasets spanning three sequencing depths (1, 10 and 100 GB). Recall generally increased with sequencing depth for all fusion callers, however the magnitude of improvement varied considerably between methods (**Figure 1C-D; Supplementary Figures 1 and 3**). *CTAT-LR-Fusion* consistently achieved the highest sensitivity but never recovered more than 30% of simulated fusion transcripts, whereas *LongGF* exhibited the lowest sensitivity across all sequencing depths. Fusion detection was very limited at 1 GB, where several callers recovered few or no simulated fusion events.

Higher sequencing depth did not consistently improve precision because additional true-positive calls were accompanied by increasing numbers of false-positive and partially reconstructed fusion transcripts **(Supplementary Figure 4)**. This trend was particularly pronounced for *FusionSeeker* and *GFSeeker*, which exhibited substantial increases in false-positive fusion calls with increasing sequencing depth.

*CTAT-LR-Fusion* and *Genion* maintained relatively low numbers of false-positive chimaeras (**Figure 1E**), but their precision declined owing to a marked increase in partially reconstructed fusion transcripts **(Supplementary Figure 5A)**. In contrast, mitochondrial, mitochondrial–genomic and self-alignment artefacts remained comparatively stable across sequencing depths, indicating that these error classes are largely independent of sequencing coverage **(Supplementary Figure 5B).**

Collectively, these findings demonstrate that deeper sequencing alone does not proportionally improve fusion detection accuracy. Instead, increasing sequencing depth amplifies both true and spurious fusion calls, highlighting the need for improved filtering strategies to maintain precision at high coverage.

#### Increasing read identity moderately improves recall but not precision

To evaluate the impact of sequencing accuracy on fusion detection, we simulated long-read datasets with mean sequence identities of 85%, 90%, and 95%, reflecting the error profiles of contemporary long-read sequencing technologies.

Increasing sequencing accuracy did not greatly improve precision or recall **(Figure 1D; Supplementary Figure 3**). *CTAT-LR-Fusion* showed the greatest gain in recall, increasing from 0.14 at 85% sequence identity to 0.28 at 95%, indicating that increased read accuracy alone is insufficient to substantially improve overall fusion detection performance. Additionally, the influence of read accuracy on false-positive fusion detection varied between callers **(Supplementary Figure 4).** *FusionSeeker* and *Genion* generated fewer false-positive fusion calls as sequence identity increased, whereas several other callers unexpectedly produced more false-positive predictions despite the higher read accuracy.

These findings demonstrate that improvements in sequencing accuracy primarily increase sensitivity but have limited impact on precision, suggesting that current computational approaches, rather than sequencing errors, are the dominant limitation for accurate long-read fusion detection.

#### CTAT-LR achieves the best overall performance, whereas JAFFAL uniquely detects tri-gene fusions

To compare the capabilities of detecting different types of fusions, we simulated 400 fusion transcripts, including 100 inter-chromosomal, 100 intra-chromosomal, 100 read-through, and 100 tri-gene fusions (**Supplementary Data 1**). Recall varied substantially between callers and fusion classes **(Figure 1E).**

*CTAT-LR-Fusion* achieved the highest overall recall (28%), performing particularly well for interchromosomal (37%), intrachromosomal fusions (50%), and read-through transcripts (26%). In contrast, *JAFFAL* was the only fusion caller to correctly identify simulated tri-gene fusions, recovering 15% of these events. Although *Genion* can report multi-gene fusions, all predicted multi-gene events were either incorrectly ordered or presented false-positive chimeras. Overall, *JAFFAL* achieved the second-highest recall across fusion classes, although *FusionSeeker* (25%) and *Genion* (17%) outperformed *JAFFAL* (15%) for read-through transcript detection **(Figure 1E).**

We next examined partially reconstructed fusion transcripts to characterise common failure modes **(Supplementary Figure 5A).** Partial predictions most frequently resulted from truncated tri-gene fusions, reversed gene order, or incorrect assignment of closely related paralogous. *FusionSeeker* generated the greatest number of partial calls, whereas over half of all predictions produced by *Genion* and *LongGF* fell into this category. In contrast, partial predictions from *JAFFAL* and *CTAT-LR-Fusion* were almost exclusively truncated tri-gene fusions, with neither method systematically reversing fusion partner order. These findings indicate that, although partial reconstruction remains common across current fusion callers, *CTAT-LR-Fusion* and *JAFFAL* produce substantially fewer structural reconstruction errors than competing methods.

#### Breakpoint localisation remains a major limitation

Accurate breakpoint localisation is essential for reconstructing fusion transcript structure. We therefore assessed the accuracy of predicted breakpoints relative to the simulated fusion junctions using a bootstrap based Two One-Sided Test (TOST) (20) with equivalence bounds of ±100 bp **(Figure 1F).** *LongGF* most accurately identified the first breakpoint, whereas *Genion* showed the smallest median error for the second breakpoint. However, both tools occasionally produced extreme breakpoint errors, including mis-localisations exceeding 100 Mb, indicating that median accuracy alone does not adequately capture reliability. Overall, *JAFFAL* provided the most consistent breakpoint predictions across both fusion junctions, whereas breakpoint localisation remained a substantial challenge for all callers **(Supplementary Figure 6).**

### Sources of false-positive fusion detection

#### False-positive fusion calls remain a major limitation across all fusion callers

False-positive fusion calls varied widely between fusion callers **(Figure 1E, Supplementary Figure 4).** *GFSeeker* generated the largest number of false-positive calls (632), followed by *FusionSeeker* (519), whereas *JAFFAL* (106) and *CTAT-LR-Fusion* (38) produced substantially fewer spurious predictions.

Some false-positive calls represented identifiable artefacts, including chimeras involving two mitochondrial genes, mitochondrial-genomic fusions and paralogous gene misalignments. These were observed primarily in *FusionSeeker*, where they accounted for 10.6% of all false-positive calls. In contrast, false-positive predictions generated by *GFSeeker*, *CTAT-LR-Fusion* and *JAFFAL* generally lacked these obvious signatures, making them more difficult to distinguish from genuine fusion transcripts.

The same pattern was observed in negative control datasets lacking simulated fusions, indicating that these artefacts arise from the computational methods rather than the simulated data. Notably, although no sense-antisense fusions were simulated *FusionSeeker* consistently reported SAS fusion transcripts, suggesting a systematic source of algorithm-specific false-positive predictions **(Supplementary Figure 5B).**

Taken together, false-positive fusion detection remains a major limitation of current long-read fusion callers, although both the frequency and characteristics of these errors differ markedly between methods.

#### Low concordance between fusion callers limits the utility of consensus filtering

Consensus fusion calling is frequently used to increase confidence in predicted fusion transcripts by retaining only events supported by multiple algorithms. However, our benchmark revealed remarkably little agreement between long-read fusion callers **(Figure 2A, Supplementary Figure 7).** Of the 191 correctly identified simulated fusions, only a single fusion was detected by all six callers, whereas 124 were supported by at least two callers. *JAFFAL* and *CTAT-LR-Fusion* each identified a substantial number of unique true-positive fusions that were not recovered by any other method, while *LongGF* detected no unique fusion events.

**Figure 2.**
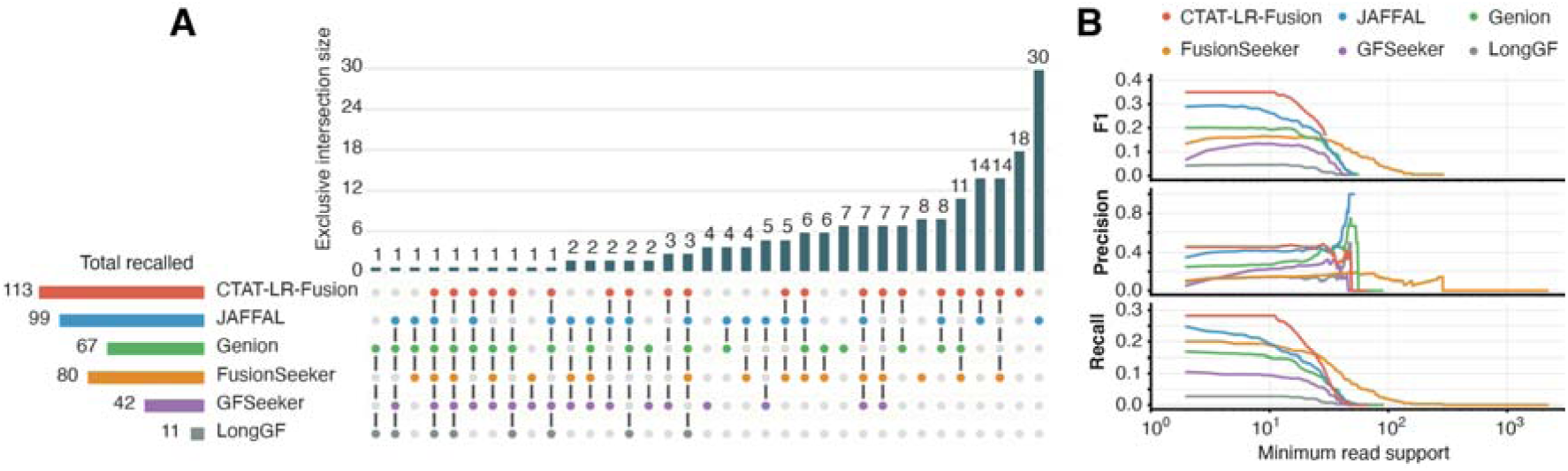
Concordance and minimum read support. **(A)** The overlap and uniqueness in recall of simulated fusions. Simulated data had 100Gb sequencing depth and 95% mean sequencing identity. **(B)** F1, Precision and Recall in relation to minimum read support. Partially recalled fusions were counted as false positives for calculation.

Although consensus filtering reduced false-positive predictions, it also eliminated many genuine fusion transcripts. Eighty-six false-positive fusions were called by at least two tools, including 12 supported by three callers, indicating that agreement between algorithms does not necessarily imply correctness. Applying increasingly stringent consensus thresholds therefore markedly reduced sensitivity, requiring agreement between four or more callers to retain only 27 true-positive fusions and 23 partially reconstructed fusion transcripts.

These findings demonstrate that consensus calling is an ineffective strategy for eliminating false-positive fusion calls and, if applied indiscriminately, substantially reduces the recovery of genuine fusion transcripts.

#### Read support filtering reveals an inherent precision-recall trade-off

Minimum read support is commonly used to filter predicted fusion transcripts, with higher thresholds assumed to improve confidence by removing lowly supported events. To evaluate this assumption, we systematically varied the minimum read support threshold for each fusion caller and quantified its effect on precision, recall, and F1 score **(Figure 2B).**

Increasing the minimum read support threshold consistently improved precision but reduced recall across all callers, demonstrating an inherent trade-off between sensitivity and specificity. The optimal threshold differed substantially between methods, with maximum F1 scores achieved between 3 and 14 supporting reads for simulations at 100 GB and 95% sequence identity. *CTAT-LR-Fusion’s* was particularly robust to increasing thresholds, maintaining its maximum F1 score up to 12 supporting reads before gradually declining. In contrast, *JAFFAL* exhibited a continuous increase in precision, ultimately reaching 100% at 47 supporting reads **(Supplementary Figure 8A-B).**

The influence of read support thresholds varied considerably between callers. *FusionSeeker* tolerated relatively stringent filtering because it assigned large numbers of supporting reads to individual fusion events. However, highly supported false-positive fusion calls were frequently retained, resulting in precision falling to zero at the highest thresholds. Similar behaviour was observed for *Genion*, *GFSeeker* and *LongGF*. In contrast, the highest-supported fusion reported by *JAFFAL* was always a genuine fusion event.

These findings demonstrate that increasing read support improves confidence only at the cost of sensitivity and that no universal threshold is suitable across fusion callers. Instead, optimal filtering strategies are caller-specific and should be selected accordingly.

### Benchmarking on cancer transcriptomes

#### Performance of long-read fusion callers in cancer cell lines

Because the complete repertoire of fusion transcripts in cancer cell lines is unknown, benchmarking real sequencing data relies on the recovery of previously validated fusion events while assessing the prevalence of obvious artefactual prediction. We therefore analysed long-read RNA sequencing datasets from the Singapore Nanopore Expression Project (SG-Nex), comprising K562 erythroleukemia and MCF7 breast cancer cell lines sequenced using ONT PCR-cDNA, direct cDNA, and direct RNA sequencing protocols (21). K562 contains six previously reported fusion transcripts, while MCF7 cells have 53 known fusions.

Consistent with the benchmark on simulated data, *CTAT-LR-Fusion* and *JAFFAL* had the highest recovery rate for known fusions across both cell lines and all library preparation protocols. *CTAT-LR-Fusion* and *FusionSeeker* identified the largest number of putative novel fusions **(Figure 3A).** However, several novel *FusionSeeker* predictions correspond to reverse-orientation calls, sense-antisense gene pairs, or antisense mappings of known fusion partners, suggesting that a proportion of these events represent algorithm-specific artefacts rather than genuine discoveries **(Figure 3B).**

**Figure 3.**
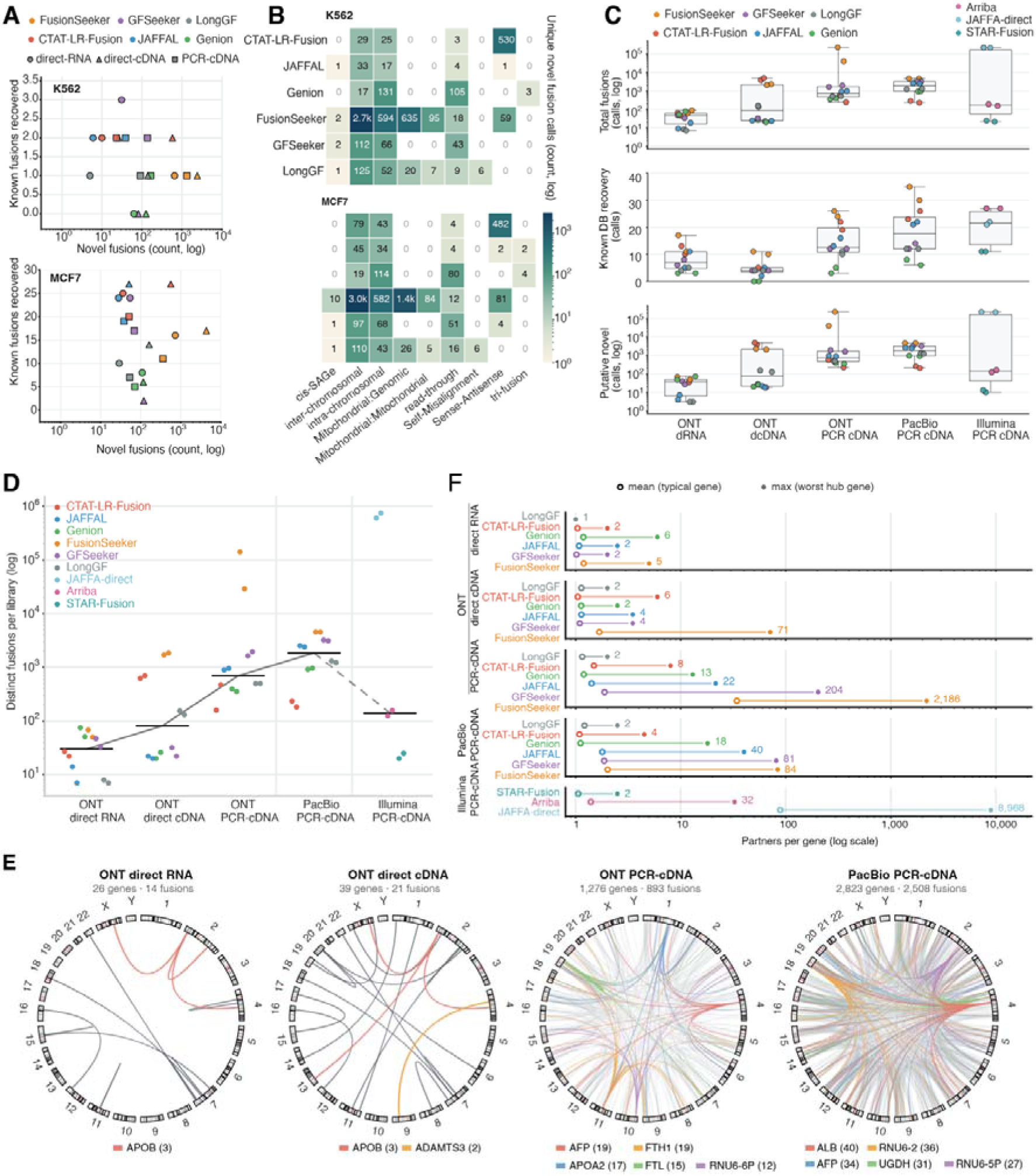
Recovery of known and putative novel fusion transcripts across sequencing platforms and fusion callers. **(A)** Recovery of known fusions in comparison to novel fusions in three types of ONT long-read RNA-seq libraries, as called by six fusion callers. **(B)** Recovery of known fusions and putative novel fusions in K562 and MCF7 cells by fusion type. **(C)** Huh7 fusion-call distribution across libraries. **(D)** Number of fusion calls reported per Huh7 library at a minimum read support of two, on a logarithmic axis, grouped by library preparation in order of increasing amplification: ONT direct RNA, ONT direct cDNA, ONT PCR-cDNA, PacBio PCR-cDNA and Illumina PCR-cDNA. Each point is one caller in one replicate library, coloured by fusion caller; the black bar marks the per-preparation median, and the grey line traces the trend between medians, dashed across the platform change to the Illumina short-read comparator. **(E)** Fusion-gene promiscuity. For each caller and library preparation (replicates averaged), a dumbbell links the mean fusion partners per gene (open circle, the typical gene) to the maximum (filled circle, the single most-connected hub gene) on a log axis; preparations run from gentlest (ONT direct RNA, top) to harshest (Illumina PCR-cDNA, bottom). **(F)** Circus plots illustrating fusion partner promiscuity (based on JAFFAL predictions). Fusion-partner networks with chromosomes drawn as the real hg38 cytoband ideogram, each arc represents a fusion transcript, with the top five hub genes coloured (others are in grey).

Both *JAFFAL* and *Genion* identified candidate tri-gene fusions, however no events were. shared between the two tools. *JAFFAL* detected tri-gene fusions exclusively in MCF7, whereas *Genion* identified several recurrent candidates in K562, including one fusion that was consistently detected across all three ONT library preparation methods.

The error profiles observed in simulated data were largely recapitulated in biological datasets. *FusionSeeker* frequently generated mitochondrial-genomic fusion artefacts, whereas *JAFFAL* and *Genion* produced no obvious false-positive chimeras. *CTAT-LR-Fusion* reported numerous sense-antisense fusion transcripts, particularly in direct-cDNA libraries. In contrast to *FusionSeeker*, these events were not observed in simulated datasets and included recurrent, highly supported fusions such as *RPL30-AS1::RPL30*, suggesting that at least a subset may represent genuine biological transcripts rather than computational artefacts.

Overall, the real-data benchmark closely mirrored the simulated analyses, confirming CTAT-LR-Fusion and JAFFAL as the strongest-performing callers while demonstrating that algorithm-specific false-positive signatures persist in biological datasets.

#### Long-read sequencing recovers more validated fusion transcripts than short-read RNA-seq

To compare long-read and short-read fusion transcript detection, we analysed Huh7 RNA sequencing data generated using multiple long-read library preparation protocols together with matched Illumina short-read RNA sequencing. Long-read datasets were analysed using the six benchmark fusion callers, whereas Illumina data were analysed using *JAFFA-direct*.

Across all datasets, 465,115 putative fusion transcripts were identified **(Figure 3C; Supplementary Figure 9).** Of these, 49 corresponded to previously reported HCC fusion transcripts, although only one had been independently validated by RT-PCR or Sanger Sequencing. Collectively, long-read sequencing recovered a greater proportion of previously reported fusion transcripts than short-read RNA sequencing, with PCR-based long-read libraries performing most consistently. PacBio Kinnex sequencing recovered the largest number of previously reported fusion transcripts (FusionSeeker), whereas Illumina and. ONT PCR-cDNA identified the greatest number of putative novel fusion events (JAFFA-direct and FusionSeeker). Notably, two fusion transcripts previously reported in HCC, but not in Huh7 cells, were detected in the Illumina data, whereas long-read libraries identified only one of these events **(Supplementary Figure 9).**

#### Long-read sequencing captures complex tri-gene fusions

One of the principal advantages of long-read RNA sequencing is its ability to recover complex fusion transcript architectures spanning multiple breakpoints. We therefore compared the detection of tri-gene fusions across sequencing platforms and library preparation methods **(Supplementary Figure 10).**

PacBio Kinnex sequencing recovered the greatest number of tri-gene fusion transcripts, followed ONT long-read libraries, whereas no tri-gene fusions were detected by in Illumina short-read data. *Genion* was the only tool to identify these putative tri-gene fusions in ONT direct-cDNA and direct-RNA libraries. Together, these findings demonstrate that long-read sequencing uniquely enables the detection of multi-segmented fusion transcripts that are inaccessible to conventional short-read RNA sequencing.

#### Detection of fusion classes varies across sequencing protocols

To determine whether sequencing protocols preferentially recover specific classes of fusion transcripts, we compared the distribution of fusion types across libraries (**Supplementary Figure 11**). PCR-based long-read libraries consistently produced mitochondrial-related chimeras, whereas only a single mitochondrial fusion was detected in the direct-RNA library. Read-through transcripts were most frequently detected by PacBio and other PCR-based libraries, while SAGe fusions were predominantly detected in Illumina datasets. *CTAT-LR-Fusion* identified substantially more sense-antisense fusion transcripts in direct-cDNA libraries than in PCR-cDNA libraries, a pattern also observed for *FusionSeeker* and in the SGNex dataset. Finally, Illumina produced by far the most predictions of putative inter- and intrachromosomal fusion transcripts compared to any long-read protocol, whereas ONT direct RNA and direct cDNA libraries consistently generated the fewest such predictions.

#### Long-read sequencing produces less promiscuous fusion landscapes

Illumina fusion transcripts were analysed using top-ranked short-read fusion callers (*Arriba*, *STAR-Fusion*, and *JAFFA-direct*) benchmarked in Haas et al. (22). Depending on the fusion caller used between 20 (STAR-Fusion) and several tens of thousands putative fusion transcripts (JAFFA-direct) were reported in short-read RNA sequencing data **(Figure 3D).** To investigate the effect of PCR amplification on fusion partner promiscuity, we modelled each dataset as a gene–gene fusion network, and quantified fusion partner promiscuity by calculating the number of distinct fusion partners associated with each gene **(Figure 3E, Supplementary Figure 12).**

Long-read datasets exhibited significantly lower fusion partner promiscuity, with genes typically participating in only two to four distinct fusion events **(Figure 3F).** In contrast, Illumina datasets showed mean fusion degrees of 187 and 227 partners per gene. Similarly, the maximum number of fusion partners observed for a single gene was orders of magnitude higher in Illumina datasets (8,828–9,510) than in long-read libraries. Among long-read protocols, ONT direct-RNA and direct cDNA sequencing showed the lowest levels of promiscuity, whereas PCR-based libraries displayed greater variability. Interestingly, fusion hub genes identified were platform specific. Unexpectedly, the high promiscuity observed in Illumina datasets was not driven by a small number of highly connected hub genes. Instead, hub dominance remained below 5%.

These findings suggest that long-read sequencing generates substantially more constrained and biologically plausible fusion landscapes than short-read RNA sequencing.

### Benchmarking long-read library preparation strategies

#### Fusion detection varies across long-read RNA-seq library preparation methods

To further investigate the influence of library preparation on fusion detection, RNA from Huh-7 cells was sequenced using ONT direct RNA, direct-cDNA and PCR-cDNA protocols together with PacBio Kinnex sequencing.

Sequencing characteristics differed substantially between library preparation methods. ONT direct-cDNA libraries consistently produced the longest reads **(Figure 4A),** whereas PacBio Kinnex generated the highest throughput (Supplementary Table 2). Read accuracy also varied between platforms, with PacBio and Illumina producing the highest-quality reads and ONT direct RNA the lowest **(Figure 4A; Supplementary Figure 13).** Gene body coverage analysis suggests that PCR-amplified libraries provide more uniform transcript coverage, whereas ONT direct sequencing protocols exhibit relatively reduced coverage toward the 31 ends of fusion transcripts **(Figure 4B).**

**Figure 4.**
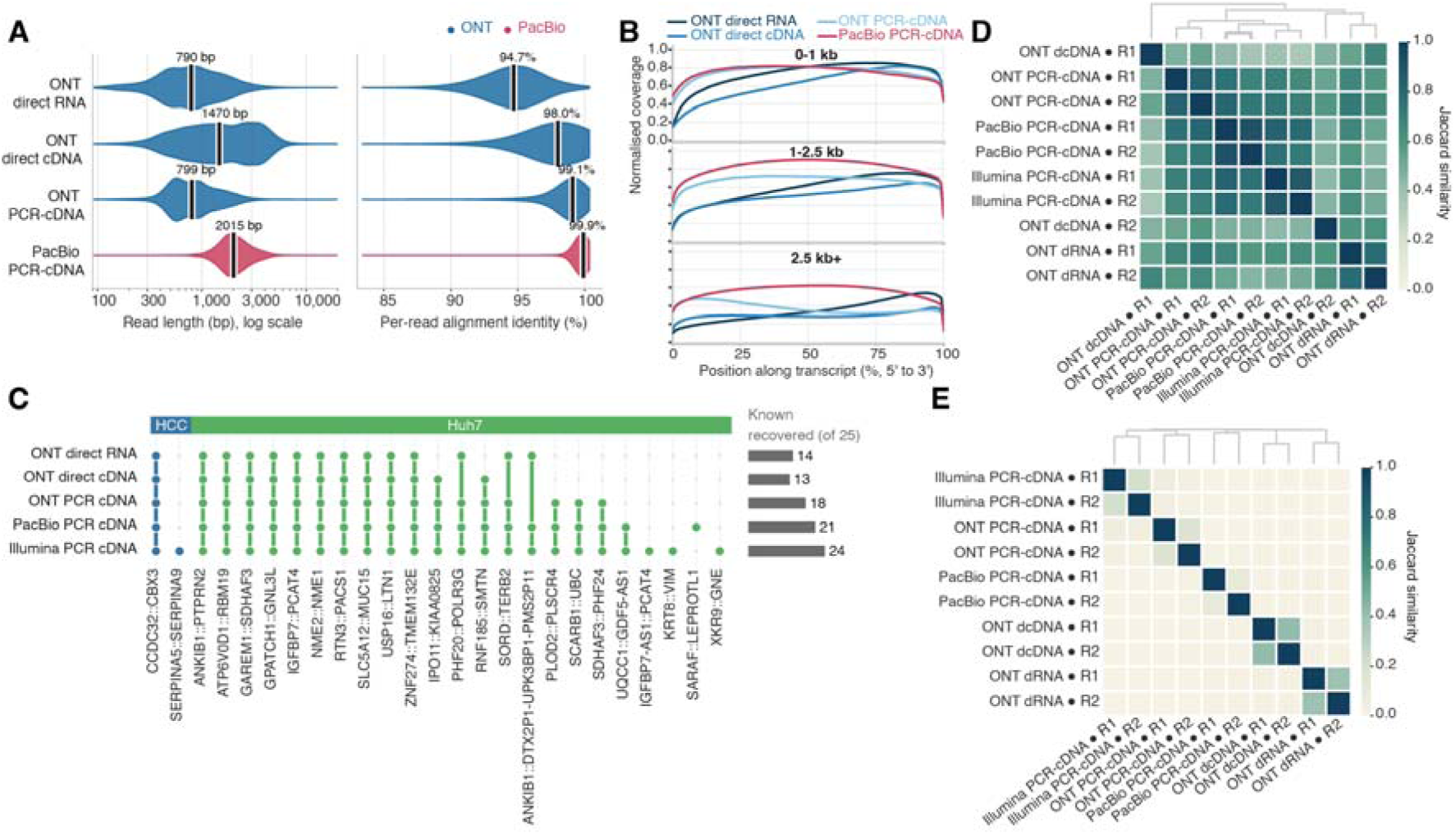
Benchmark long-read library preparation strategies. **(A)** Read length and identity by library preparation. Distributions of per-read length (left, logarithmic axis) and per-read alignment identity (right) for the four long-read Huh7 library preparations: ONT direct RNA, ONT direct cDNA, ONT PCR-cDNA and PacBio PCR-cDNA. Each preparation is shown as a horizontal violin coloured by sequencing platform (ONT, PacBio), pooling its two replicates over all primary alignments per library. Alignment identity is defined as (aligned bases minus edit distance) divided by aligned bases, with introns and soft or hard clips excluded so that spliced alignments are not penalised. Medians are annotated on each violin: read lengths of 790, 1,470, 799 and 2,015 bp, and identities of 94.7, 98.0, 99.1 and 99.9 per cent, respectively. **(B)** Coverage across the transcript body for the four long-read Huh7 libraries, faceted by transcript length (0-1 kb, 1-2.5 kb, 2.5 kb+). The x-axis denotes the position along the transcript (0% = 5⍰ end, 100% = 3⍰ end); the y-axis represents the normalised coverage. Coverage was computed from all primary alignments to the GENCODE transcriptome: for every transcript with at least 10 reads, the fraction of its reads covering each percentile position was calculated, and these per-transcript profiles were then averaged with equal weight within each length bin (the standard gene-body-coverage approach, so that highly expressed transcripts do not dominate); the two replicates were pooled. Lines are coloured by platform, the three ONT protocols as shades of blue and PacBio in red. **(C)** Recovery of 25 curated known Huh7 or HCC fusions across library preparations detected by at least one library. Bar chart at right indicates the total number recovered per library. **(D)** Each library replicate is represented by its set of distinct fusions, defined as canonical Ensembl gene-identifier pairs so that A::B and B::A are counted once. Only calls flagged as known fusions are included, retained at a minimum of two spanning reads or read pairs. **(E)** Pairwise Jaccard similarity of putative novel fusion calls between library replicates. In C-E, A combination of results from all six benchmarked long-read fusion callers was used for the long-read data, and from all three short-read callers (JAFFA-direct, Arriba and STAR-Fusion) for the Illumina short-read data.

These protocol-specific differences directly influence fusion detection performance. As demonstrated in the simulated benchmark, longer reads, higher sequencing depth and increased read accuracy each improve the recovery of fusion transcripts, although these gains are accompanied by distinct trade-offs in precision and false-positive detection. Consequently, differences in library preparation should be considered when comparing fusion landscapes across studies.

#### PCR-cDNA libraries provide the most consistent recovery of validated fusion transcripts

To assess the ability of different protocols to recover previously reported fusion transcripts, we curated a reference set of known Huh7 and HCC fusion events from the literature and public databases **(Figure 4C, Supplementary Data 3).**

Of the 34 previously reported Huh7 fusion transcripts, 21 were recovered across the evaluated sequencing libraries. PacBio Kinnex recovered the greatest number of known fusions (20), followed by ONT PCR-cDNA (17), whereas Illumina recovered 15. Interestingly, none of the three Huh7 fusion transcripts previously validated by RT-PCR or Sanger sequencing (*MAN2A1::FER, CCNH::C5orf30,* and *SLC45A2::AMACR*) were detected in our datasets (23, 24). In contrast, the experimentally validated HCC fusion *C15orf57::CBX3* was recovered by all sequencing protocols (25).

To assess reproducibility, we compared biological replicates using the Jaccard similarity of recovered known fusion transcripts **(Figure 4D).** PacBio libraries exhibited the highest within-platform similarity (Jaccard = 0.91), exceeding that of Illumina (0.81), whereas ONT PCR-cDNA and ONT’s direct cDNA showed the lowest similarity (0.40).

Together, these results demonstrate that PCR-based long-read sequencing, particularly PacBio Kinnex, provides the most reliable recovery of previously characterised fusion transcripts.

#### Novel fusion discovery remains highly protocol-dependent

We next compared the reproducibility of putative novel fusion transcript discovery **(Figure 4E).** In contrast to known fusion recovery, Jaccard similarity between biological replicates declined markedly across all sequencing protocols. PCR-based libraries exhibited the greatest drop, with within-platform similarities falling to 0.21 for Illumina, 0.12 for PacBio and 0.09 for ONT PCR-cDNA. ONT direct RNA and direct-cDNA retained comparatively higher reproducibility (0.40–0.49), although similarity between different library preparation methods remained low.

These observations indicate that novel fusion discovery remains highly dependent on sequencing protocol and library preparation, suggesting that many putative fusion transcripts are not reproducibly detected across independent experiments.

## Discussion

The performance of transcriptome-wide fusion discovery is shaped by four interdependent components, namely sequencing technology, library preparation, computational algorithms, and analysis strategies. Accurate detection of fusion transcripts underpins studies of cancer, inherited disease, and evolutionary biology, and is increasingly important for precision medicine. Although long-read RNA sequencing has transformed the ability to resolve full-length fusion transcripts and complex multi-segmented fusion architectures, the relative performance of available computational tools and sequencing strategies has remained largely uncharacterised. Such evaluation is crucial for downstream analyses and biological interpretation (21, 26). Here, we present the first comprehensive benchmark of long-read fusion detection workflows across multiple fusion callers, sequencing platforms, and library preparation methods. Our analyses revealed that current fusion callers recover fewer than one-third of simulated fusion transcripts, breakpoint localisation remains a major limitation, and false-positive fusion calls are pervasive across all methods. We further showed that sequencing depth, read accuracy, consensus filtering, and minimum read-support thresholds each involve substantial trade-offs. Of note, ONT PCR-cDNA combined with *CTAT-LR-Fusion* provided the best overall balance between sensitivity and precision and *JAFFAL* uniquely enables reliable detection of tri-gene fusions **(Table 1).** Together, these findings establish practical guidelines for long-read fusion transcript discovery while highlighting key methodological challenges that remain to be addressed.

**Table 1.**
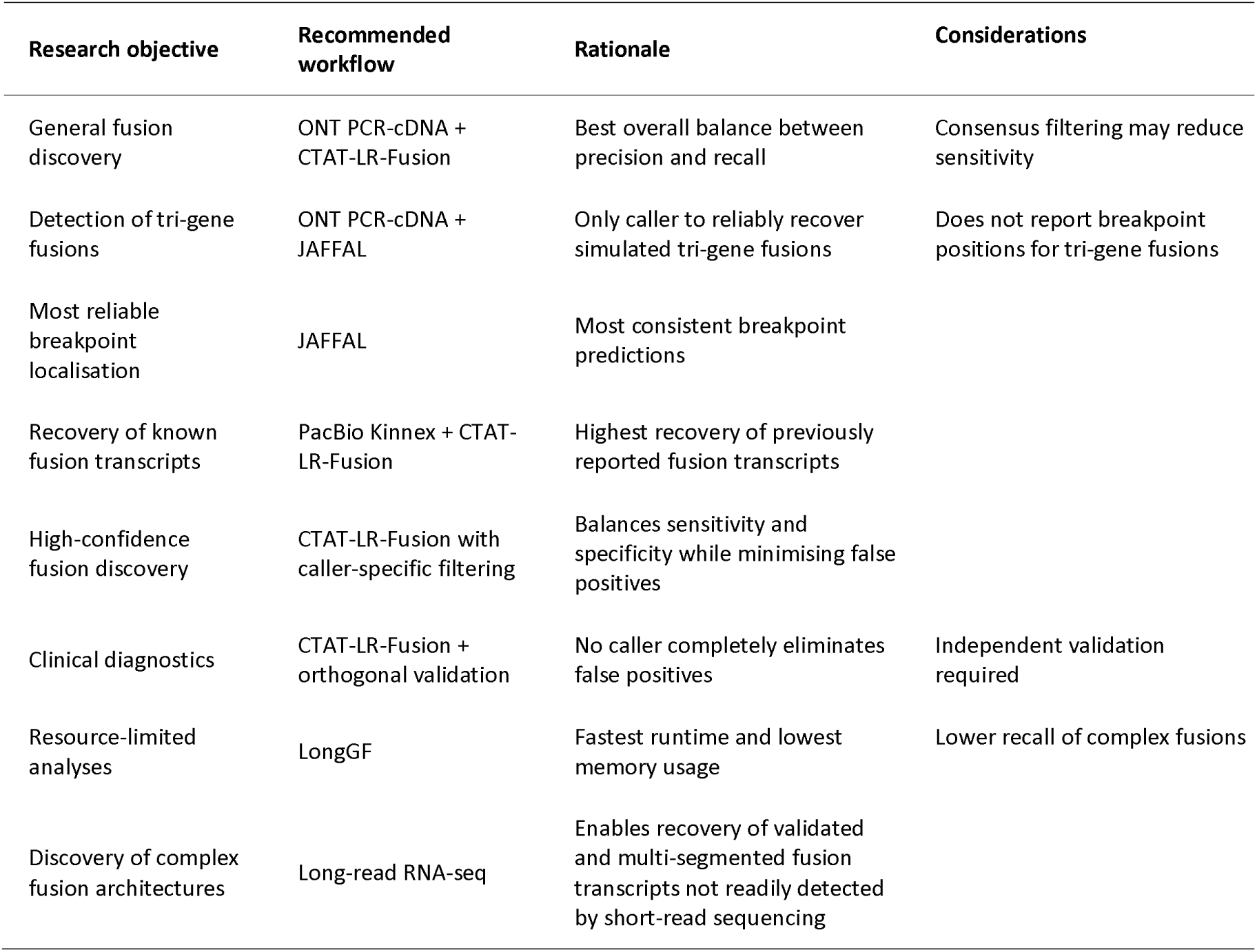
Recommended fusion transcript discovery workflows.

| Research objective | Recommended workflow | Rationale | Considerations |
| --- | --- | --- | --- |
| General fusion discovery | ONT PCR-cDNA + CTAT-LR-Fusion | Best overall balance between precision and recall | Consensus filtering may reduce sensitivity |
| Detection of tri-gene fusions | ONT PCR-cDNA + JAFFAL | Only caller to reliably recover simulated tri-gene fusions | Does not report breakpoint positions for tri-gene fusions |
| Most reliable breakpoint localisation | JAFFAL | Most consistent breakpoint predictions |  |
| Recovery of known fusion transcripts | PacBio Kinnex + CTAT-LR-Fusion | Highest recovery of previously reported fusion transcripts |  |
| High-confidence fusion discovery | CTAT-LR-Fusion with caller-specific filtering | Balances sensitivity and specificity while minimising false positives |  |
| Clinical diagnostics | CTAT-LR-Fusion + orthogonal validation | No caller completely eliminates false positives | Independent validation required |
| Resource-limited analyses | LongGF | Fastest runtime and lowest memory usage | Lower recall of complex fusions |
| Discovery of complex fusion architectures | Long-read RNA-seq | Enables recovery of validated and multi-segmented fusion transcripts not readily detected by short-read sequencing |  |

### The best tools for the job

Our benchmark demonstrates that no single fusion caller is optimal across all performance metrics, reflecting the diverse algorithmic strategies used for long-read fusion detection. Nevertheless, clear strengths emerged. *CTAT-LR-Fusion* consistently achieved the best overall balance between sensitivity and precision, whereas *JAFFAL* uniquely enabled reliable detection of complex tri-gene fusions and produced the most consistent breakpoint localisation. These findings were reproduced across both simulated datasets and cancer cell lines, indicating that their superior performance generalises across sequencing protocols and biological datasets.

The superior performance of *CTAT-LR-Fusion* and *JAFFAL* likely reflects their shared use of two-stage alignment strategies, in which candidate chimeric reads undergo additional realignment and breakpoint refinement before fusion calling. This contrasts with methods that rely primarily on single-pass split-read alignments, which were generally faster but exhibited substantially higher false-positive rates or lower recall. Although *CTAT-LR-Fusion* achieved the highest overall recall, no caller recovered more than 30% of simulated fusion transcripts, indicating that accurate fusion detection remains a major computational challenge rather than a limitation of sequencing technology alone.

The evaluated callers also differed considerably in the complexity of fusion structures they could reconstruct. *JAFFAL* was the only method to reliably recover simulated tri-gene fusions and was the only caller that consistently preserved the correct order of fusion partners. In contrast, several callers frequently generated partial reconstructions or reversed fusion orientations, highlighting that reconstructing complex transcript architectures remains considerably more challenging than identifying canonical two-gene fusions.

From a practical perspective, the optimal fusion caller depends on the intended application. For general fusion discovery, *CTAT-LR-Fusion* currently provides the most robust balance between sensitivity and precision. Studies focusing on complex fusion architecture, including tri-gene fusions, would benefit from *JAFFAL*, whereas *LongGF* remains attractive for computationally constrained analyses because of its low memory requirements and rapid execution. *FusionSeeker* offers fast exploratory analyses but requires careful interpretation because of its relatively high false-positive rate (Table 1).

Finally, the utility of downstream analyses depends not only on whether a fusion is detected but also on the information provided by the fusion caller. *JAFFAL* reports strand-aware breakpoint annotations for canonical fusion transcripts, whereas *Genion* provides genomic intervals for each fusion partner without strand information. *FusionSeeker* and *LongGF* report breakpoint positions but comparatively limited genomic context. As fusion transcript analyses increasingly extend beyond simple detection towards functional interpretation, richer breakpoint annotation and more accurate reconstruction of complex transcript structures will become increasingly important.

### Study limitations

Several limitations should be considered when interpreting our benchmark. First, although simulated datasets enabled objective assessment of precision, recall and breakpoint accuracy, they cannot fully capture the complexity of biological transcriptomes, including variable transcript abundance, mapping ambiguity, and uncharacterised fusion architectures. We therefore complemented the simulations with publicly available cancer cell line datasets and newly generated Huh7 sequencing data, although the complete repertoire of true fusion transcripts in these models remains unknown. Second, our library comparison relied primarily on previously reported fusion transcripts rather than independent experimental validation. Although we curated validated fusion events from the literature, additional RT-PCR or long-range sequencing would be required to confirm the biological validity of newly identified fusion transcripts, particularly complex tri-gene fusions. Finally, although this benchmark includes the major published long-read fusion callers available at the time of analysis, some emerging tools could not be evaluated. *AERON* (*27*) failed to execute reproducibly despite extensive troubleshooting, while *FLAIR-Fusion* (*28*) had not yet undergone peer review when this study commenced. As the field continues to evolve, future benchmarks should incorporate newly developed algorithms as they become sufficiently mature.

### Future Directions

Our benchmark highlights several opportunities for improving long-read fusion transcript detection. Despite rapid advances in sequencing technology, computational methods remain the principal limitation, with no evaluated caller recovering more than one-third of simulated fusion transcripts. Future algorithm development should therefore prioritise improved breakpoint localisation, reconstruction of complex multi-segmented fusion transcripts, and reduction of biologically plausible false-positive predictions. Future fusion callers should shift from identifying individual breakpoints towards reconstructing complete fusion transcript architectures.

Our findings also question the widespread use of consensus fusion calling as a universal confidence filter. Although agreement between callers reduced some false-positive predictions, it also removed a substantial proportion of genuine fusion transcripts because current algorithms exhibit surprisingly limited concordance. Future consensus approaches should therefore incorporate caller-specific confidence estimates or probabilistic frameworks rather than simple voting strategies. Recently proposed methods such as *GFVoter* (29) represent promising steps in this direction, although our results suggest that consensus alone is unlikely to overcome the current limitations of long-read fusion detection.

Finally, as long-read sequencing continues to improve in throughput and accuracy, benchmarking efforts should extend beyond canonical gene fusions to encompass complex transcript architectures, including tri-gene and higher-order fusion transcripts, fusion isoforms, and their functional consequences. Establishing experimentally validated reference datasets will be essential for enabling the next generation of benchmarking studies.

Collectively, our benchmark demonstrates that long-read RNA sequencing has matured into a powerful technology for fusion transcript discovery, but current computational methods remain the primary bottleneck. Rather than simply increasing sequencing depth or read accuracy, future advances will depend on algorithms capable of accurately reconstructing complete fusion transcript architectures while reducing biologically plausible false-positive predictions. As long-read sequencing continues to improve, experimentally validated benchmark datasets and more sophisticated computational approaches will be essential for enabling reliable fusion discovery in both biological research and clinical diagnostics.

## Methods

We established a comprehensive benchmarking framework for transcriptome-wide fusion discovery using long-read RNA sequencing **(Figure 1A; Supplementary Figure 13).** The benchmark combined simulated datasets with experimentally derived transcriptomes from three cancer cell lines and evaluated six long-read fusion callers across multiple sequencing platforms, library preparation methods, and analysis strategies.

### Datasets

An overview of the benchmarking workflow, including simulated and experimental datasets, sequencing platforms, preprocessing, alignment, and fusion-calling pipelines, is provided in **Supplementary Figure 14.**

#### Simulation datasets

Fusion transcripts were simulated using *Fusim* (30) with the GRCh38 (hg38) reference genome and GENCODE v43 gene annotation. Five classes of fusion transcripts were generated (100 per class): inter-chromosomal fusions, intra-chromosomal fusions, read-through transcripts, tri-gene fusions, and self-fusions (**Supplementary Data 1**).

Preliminary analyses showed that the simulated self-fusions were not detected by any evaluated fusion caller, likely because they more closely resemble circular RNA species than the sense–antisense or gene duplication events targeted by current long-read fusion detection algorithms. Consequently, self-fusions were excluded from precision and recall calculations, resulting in a benchmark comprising 400 simulated fusion transcripts.

Long-read RNA sequencing datasets were simulated using *Badread* (v04.1) (31) based on the GENCODE human reference transcriptome. To generate biologically realistic transcript abundance distributions, transcript-specific sequencing depths were assigned by appending `depth = <value>` to FASTA headers. Values were assigned based on transcript quantification data from a HepG2 ONT direct-RNA sequencing dataset (ENCODE accession ENCFF382KCL), with only transcripts annotated as *Known* assigned measured abundances. Transcript abundances were derived from an independent HepG2 long-read RNA sequencing dataset to generate biologically realistic, non-uniform transcript expression distributions. All other transcripts were assigned a depth of 1.0. Simulated fusion transcripts inherited the abundance of their respective head gene.

Datasets were generated using the *Badread* Nanopore2023 error model at three sequencing depths (1Gb, 10Gb, 100Gb) and three mean read identities (85%, 90% and 95%). This resulted in nine simulated sequencing scenarios spanning combinations of sequencing depth and read identity. For all simulations, the maximum read identity was fixed at 99.5% with a standard deviation of 2.5%.

Simulation parameters were otherwise based on the default *Badread* configuration. Adapter sequences were retained, whereas junk reads, random reads and glitch reads were disabled. Artificial chimeric reads were simulated at a rate of 0.1%. Simulations of the 1 and 10 Gb datasets used a random seed of 12. Because of computational resource constraints, the 100 Gb datasets were generated by concatenating ten independent 10 Gb simulations using random seeds 1-9 and 12.

#### Public long-read datasets

Long-read RNA sequencing datasets from the K562 erythroleukaemia and MCF7 breast cancer cell lines were obtained from the Singapore Nanopore Expression Project (SGNex) (21). ONT sequencing data comprised PCR-cDNA (unstranded), direct cDNA and direct RNA libraries. For each cell line and library, FASTQ files from individual sequencing runs were merged and subsequently down-sampled using *rasusa* to generate datasets spanning sequencing depths of 1, 2.5, 5, 7.5 and 10 Gb, where sufficient sequencing data were available. These datasets were used to evaluate the effect of sequencing depth on fusion transcript detection in experimentally derived long-read RNA sequencing data.

#### Cell Culture

Huh7 hepatocellular carcinoma cells were cultured in high-glucose Dulbecco’s Modified Eagle Medium (DMEM, Gibco™, 10566016) supplemented with GlutaMAX, 10% heat-inactivated foetal bovine serum (FBS), and 1% penicillin-streptomycin. Cells were maintained at 37°C in a humidified incubator with 5% CO_2_ and harvested for RNA extraction at approximately 70% confluency.

#### RNA Extraction

Total RNA was extracted with the RNeasy Plus Mini Kit (Qiagen, Cat No. 74134) following the manufacturer’s instructions. RNA purity was assessed using a NanoDrop spectrophotometer (A260/280 and A260/230 ratios), RNA concentration was determined using the Qubit™ RNA Broad Range (BR) Assay kit (Thermo Fisher Scientific, Q10210), and integrity assessed using RNA ScreenTape on the Agilent TapeStation system (Agilent Technologies).

#### Library preparation and RNA sequencing

For Huh7 cells, sequencing libraries were prepared using ONT direct-RNA (SQK-RNA004), direct-cDNA (SQK-LSK114), and PCR-cDNA (SQK-PCS114) library preparation kits. Two biological replicates were generated for each library preparation method. Additional full-length RNA sequencing was performed using the PacBio Kinnex workflow, and matched short-read RNA sequencing was performed on an Illumina NovaSeq platform (150-bp paired-end). ONT libraries were sequenced on individual MinION flow cells (R.10.4 chemistry for cDNA libraries and RNA004 chemistry for direct RNA libraries) until flow-cell exhaustion. Basecalling was performed using *dorado* (v7.4.12) through *MinKNOW* (v24.06.10). PacBio Kinnex and Illumina NovaSeq libraries were prepared and sequenced by the Australian Genome Research Facility (AGRF) according to the manufacturer’s recommended protocols.

#### Read processing

Illumina paired-end reads were processed using fastp (v.0.23.4), including adapter trimming, polyG trimming, removal of unpaired reads, and filtering reads shorter than 36bp. PacBio Kinnex data were processed using the IsoSeq3 clustering workflow (cluster2). Consensus transcript reads were exported from the resulting transcripts.bam file in FASTQ format using *bam2fastq.*

ONT PCR-cDNA and direct-cDNA libraries were basecalled using *Dorado* (v7.3.11), whereas direct-RNA libraries were basecalled using Dorado (v7.4.12). Super-accurate basecalling models were used throughput, and adapter trimming was performed using Dorado. Adapter sequences were removed from simulated reads using *PorechopABI* (v0.5.0) (32). Sequencing quality was assessed for both simulated and experimental datasets using *FastQC* (v12.0) (33), and quality reports were summarised with *MultiQC* (34). Reads were aligned to the hg38 reference genome using minimap2 (v2.28) (35). Although *Genion* recommends *desalt* as its default aligner, *minimap2* was used for all evaluated fusion callers to ensure a consistent benchmarking framework. *Genion* accepts alignments generated by any splice-aware long-read aligner provided they are supplied in PAF format. SAM files were converted to BAM, sorted and indexed using *SAMtools* (v1.20). Read counts following adapter trimming are summarised in **Supplementary Table 1.**

### Benchmark workflow

#### Reference genomes

The hg38 reference genome and corresponding GENCODE v44 gene annotation were used throughout the benchmark unless otherwise require by individual fusion callers. Because *JAFFAL* currently required UCSC-compatible annotations, the corresponding GENVODE v43 annotation obtained from the UCSC Genome Browser was used. *Genion* analyses were performed using Ensembl release 110 annotations, which corresponds to GENCODE v44.

*Genion* additionally requires reference files describing transcript sequence similarity and segmental duplications. These were generated following the *Genion* documentation using all-to-all alignment of the Ensembl cDNA reference transcriptome and the hg38 segmental duplication annotation downloaded from the UCSC Genome Browser (ftp://hgdownload.soe.ucsc.edu/goldenPath/hg38/database/genomicSuperDups.txt.gz).

#### Fusion caller configuration

Because the evaluated fusion callers differ in their default configurations, we standardised reference genomes, annotations, alignment software and reporting formats wherever possible to minimise methodological bias unrelated to fusion detection performance. For *JAFFAL*, the bundled reference genome was replaced with the UCSC equivalent of GENCODE v43 and the bundled versions of *minimap2* and *SAMtools* were updated to v2.28 and v1.20, respectively. *JAFFAL’s* optional setting to retain read-through transcripts was enabled to ensure consistent evaluation across all simulated fusion classes. Several fusion callers report partners using gene symbols rather than Ensembl gene IDs. To enable direct comparison between callers and simulated truth sets, gene symbols were converted to Ensembl gene IDs using *biomaRt*, while simulated Ensembl gene IDs were mapped back to gene symbols where required. Unless otherwise stated, only fusion calls supported by at least two sequencing reads were retained for downstream analyses.

#### Benchmark evaluation criteria

To enable direct comparison between fusion callers, all fusion predictions were standardised to Ensembl gene identifiers, which provide more stable identifiers than gene symbols across annotation releases. Gene symbols reported by individual fusion callers were converted to Ensembl gene IDs using *biomaRt*. Fusion predictions were considered correct only when both partner genes matched the simulated truth set. For paralogous genes, including homologues located on chromosomes X and Y, the correct chromosomal origin was also required. For example, simulated *AKAP17A::ZBED1* fusions located on chromosome Y were not considered correctly identified when reported using the homologous chromosome X loci. Some flexibility was permitted for genes represented by multiple equivalent annotations or overlapping loci. For example, *TXLNGY* is represented by overlapping Ensembl gene identifiers (ENSG00000131002 and ENSG00000291033), and equivalent annotations were accepted during benchmarking where appropriate.

##### True positives (TP)

Fusion calls that correctly identify a simulated or validated reference fusion. A call is considered a true positive if it matches the reference fusion according to the predefined matching criteria (e.g. gene partners and directionality).

##### False positives (FP)

Fusion calls that do not correspond to any reference fusion. These include spurious fusion predictions, incorrect gene-pair assignments, and other calls failing the matching criteria.

##### False negatives (FN)

Reference fusions that were not detected by the caller.

##### Partial recalls (PR)

Reference fusions that were only partially recovered. In this benchmark, a partial recall denotes a call in which fusion partners are correctly identified but the complete reference fusion is not recovered (e.g. incorrect chromosome assignment for sex paralogues, an incomplete multi-gene fusion, reverse or incorrect ordering of partner genes, incorrect Ensembl ID, or another predefined partial match).

##### Recall (sensitivity)

The proportion of reference fusions that were successfully detected.

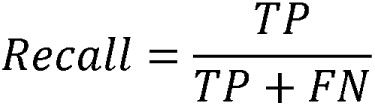

##### Precision

The proportion of reported fusion calls that are correct.

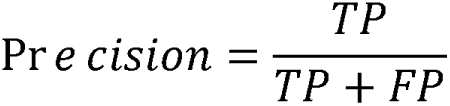

##### F1 score

The harmonic mean of precision and recall.

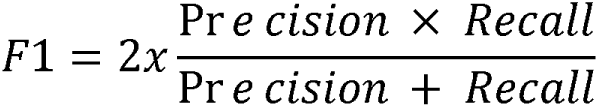

#### Fusion-call similarity between libraries

Pairwise similarity of fusion calls between Huh7 libraries was computed from the aggregated fusion call table. For each replicate, fusion calls were filtered to a minimum support of two spanning reads (long-read libraries) or two spanning pairs (Illumina). The fusion set for each replicate was then defined as the union of retained calls across all fusion callers applied to that library. Each fusion was reduced to a canonical, unordered gene-pair identifier by extracting the two Ensembl gene identifiers and sorting them alphabetically, such that reciprocal orientations (A::B and B::A) were treated as the same fusion event. Each replicate was therefore represented as a set of unique gene pairs. Three analyses were performed by filtering calls according to their annotation category before set construction: known fusions (categories *Known*, *Reverse Known*, *contains Known fusion between genes 1 and 2*, and *contains Known fusion between genes 2 and 3*), putative novel fusions (*Putative Novel*), and all fusions (no category filter). Similarity between replicate sets A and B was quantified using the Jaccard index,

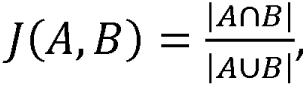

which was calculated for every pair of replicates to generate a symmetric 10 × 10 similarity matrix. Self-similarity was defined as 1, while comparisons involving an empty union were assigned a value of 0. For visualization, the similarity matrix was converted to a distance matrix (*D = 1 - J*) and hierarchically clustered using average linkage (UPGMA; scipy.cluster.hierarchy.linkage(method="average")). The resulting dendrogram determined the row and column order of the heatmap but did not modify the underlying similarity values.

#### Statistical data analysis

All statistical tests were performed in R v4.4.2 and RStudio (v2023.6.1.524). The TOSTER v0.8.6 R package was used for the bootstrap-based TOST (36).

## Declarations

### Ethics approval and consent to participate

Not applicable

### Consent for publication

Not applicable

### Availability of data and materials

The datasets generated and/or analysed in this study are available in Gene Expression Omnibus (GEO) and the Sequencing Read Archive (SRA) under accession: XXXXXXXXX. SGNex ONT cell line sequencing is available at https://github.com/GoekeLab/sg-nex-data (21). The benchmarking documentation and code are made available through GitHub: https://github.com/RDorney/LRFusionBenchmark

### Competing interests

The authors declare that they have no competing interests.

### Funding

This work was supported by the National Health and Medical Research Council (Grant #1196405 to U.S.); the Tropical Australian Academic Health Centre (Grant SF01124 to U.S.); the Townsville University Hospital (Grants #THHSSERTA RPG05 2024, #THHSSERTA RPG15 2024, #THHSSERTA RCG05 2024, and #RPG09_2025 to U.S.); Tropical Australian Academic Health Centre Limited Research Seed Grant (SF000121 to L.H.), Townsville Hospital Health Service-Study Education Research Trust Account, Project and Capacity Building Grants (RPG1 2023 and RCG2 2023 to U.S. and L.H.), and the Centre for Bioinformatics and Molecular Biology (seed grant to R.D.).

### Authors’ contributions

R.D. and S.W. contributed equally to this work. R.D. and U.S. conceived of ideas in this manuscript and designed the research. R.D. and S.W. performed the analysis of data and generated the figures. R.D. wrote the preliminary draft of the manuscript. J.H. supported the setup of Oxford Nanopore sequencing in the lab and helped experimental planning and analysis. L.H. provided the liver cancer cells. Both U.S. and L.H. supervised the work and critically reviewed the manuscript. All authors have read and approved the final version of the manuscript.

## Acknowledgements

Not applicable

## Supplementary Materials

**Supplementary Table 1:**
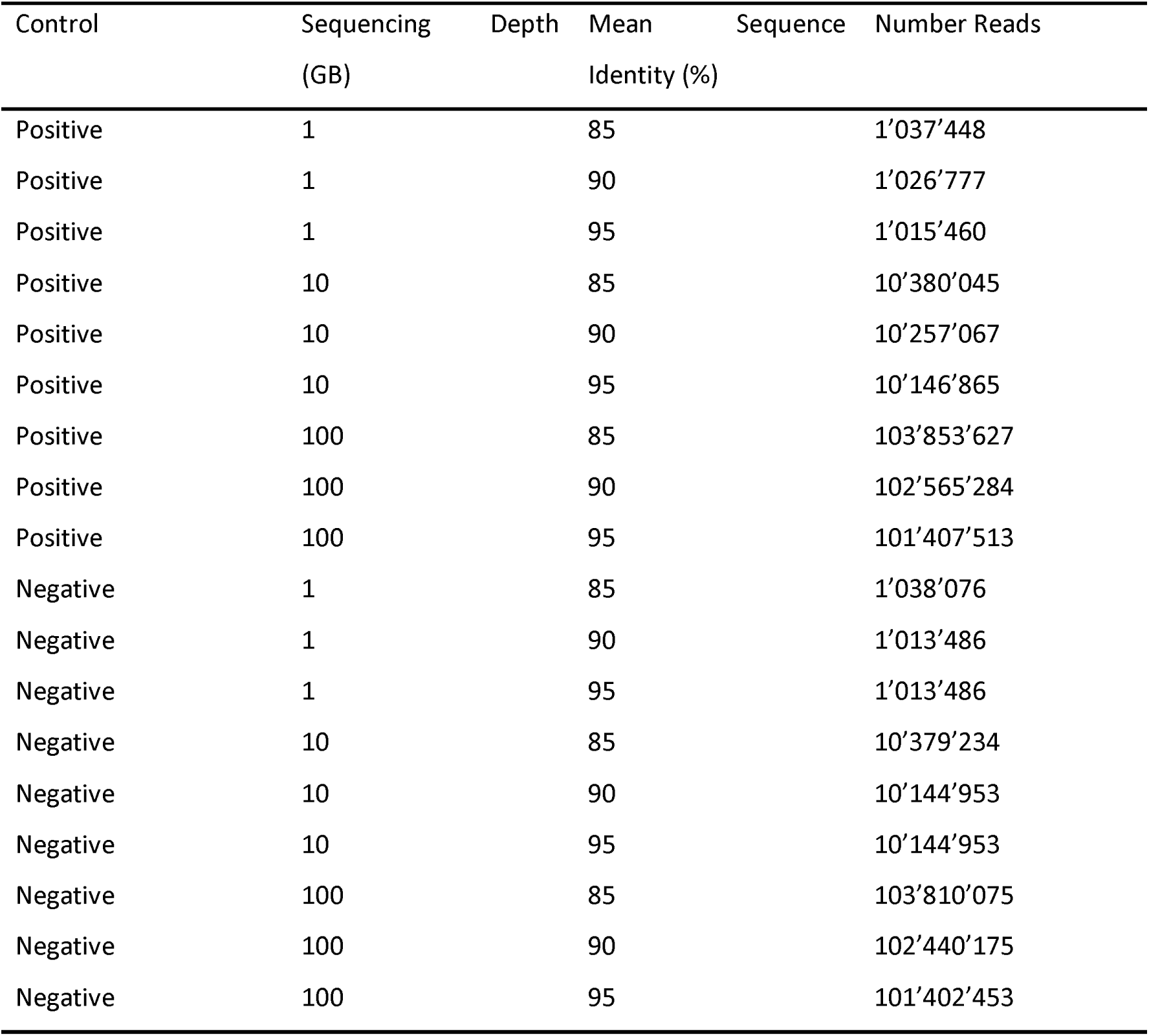
Number of reads in each simulated file following processing with Porechop ABI.

**Supplementary Table 2:** Key library performance metrics.

| Library<br>prep | # Reads |  | Yield (Gb) |  | N50 (bp) |  | Identity (%) |  |
| --- | --- | --- | --- | --- | --- | --- | --- | --- |
|  | R1 | R2 | R1 | R2 | R1 | R2 | R1 | R2 |
| ONT direct-RNA | 5,731,969 | 3,837,353 | 5.67 | 3.66 | 1,349 | 1,334 | 94.9 | 94.5 |
| ONT direct-cDNA | 3,625,384 | 7,092,066 | 5.81 | 14.52 | 2,515 | 2,954 | 98.4 | 97.9 |
| ONT PCR-cDNA | 17,673,844 | 18,109,856 | 16.65 | 17.76 | 1,038 | 1,124 | 99.1 | 99.1 |
| PacBio<br>Kinnex | 25,837,052 | 25,422,863 | 56.64 | 54.66 | 2,274 | 2,228 | 99.9 | 99.9 |

**Supplementary Figure 1.**
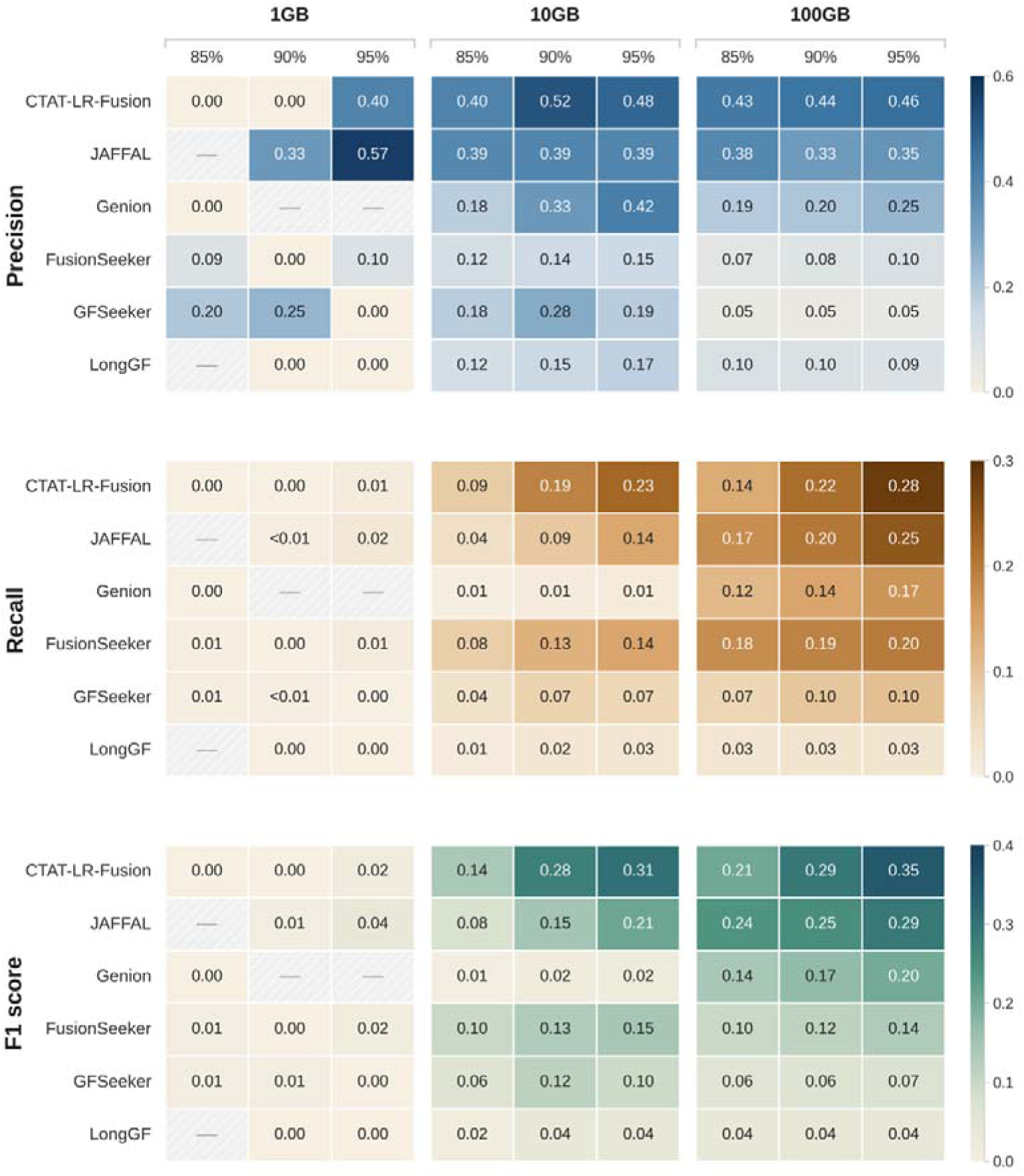
Precision, recall and F1 scores of six long-read RNA-seq fusion callers across simulated sequencing depths and sequence identities.

**Supplementary Figure 2.**
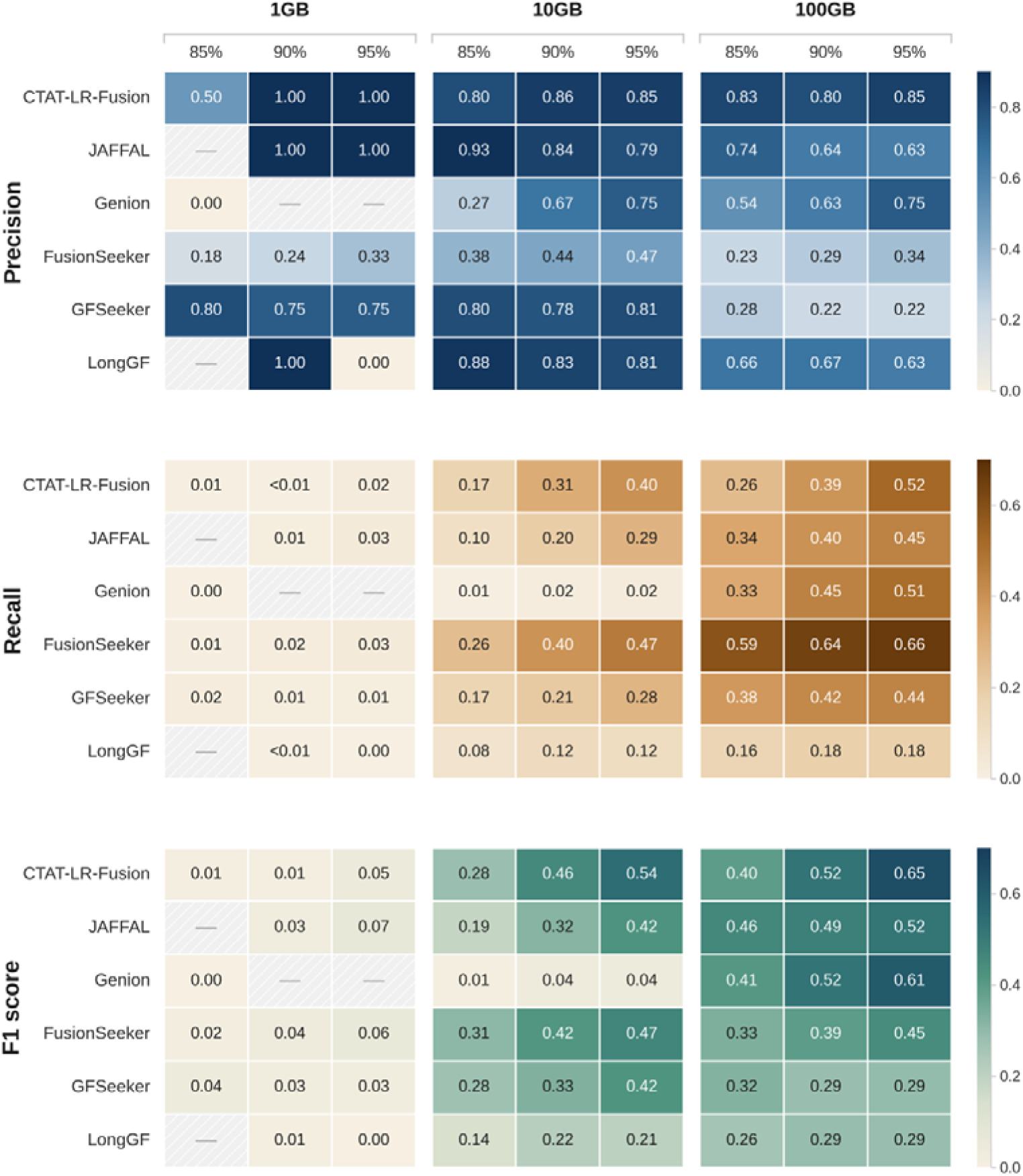
Recalculated precision, recall and F1 scores of six long-read RNA-seq fusion callers across simulated sequencing depths and sequence identities, where partial calls are counted as true recall.

**Supplementary Figure 3.**
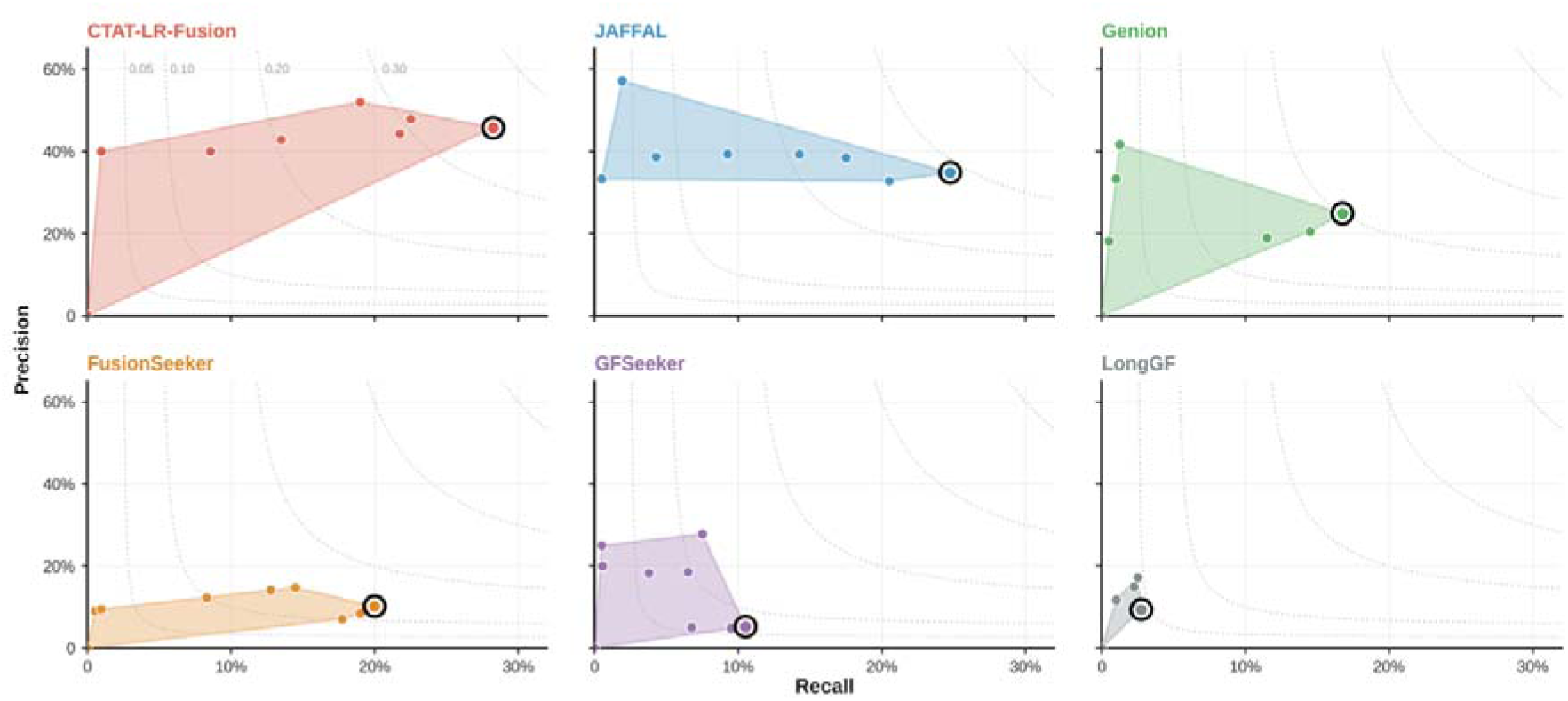
Recall vs precision of fusion callers across simulated sequencing depths and mean sequence identities. Partially recalled fusions were counted as false positives for calculation.

**Supplementary Figure 4.**
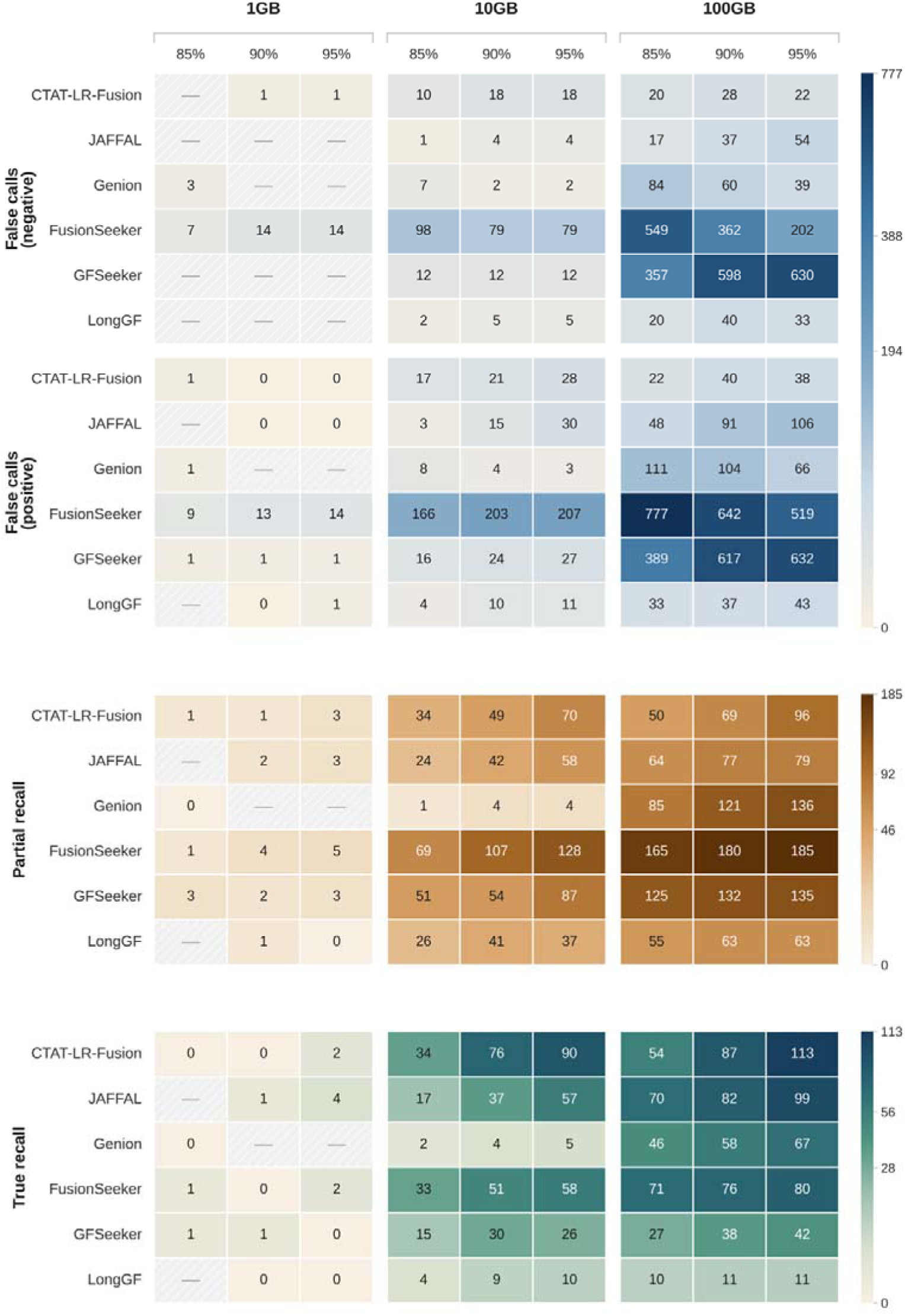
Fusion caller false calls, partially-recalled, and true fusions for different sequencing depths and sequence identities.

**Supplementary Figure 5.**
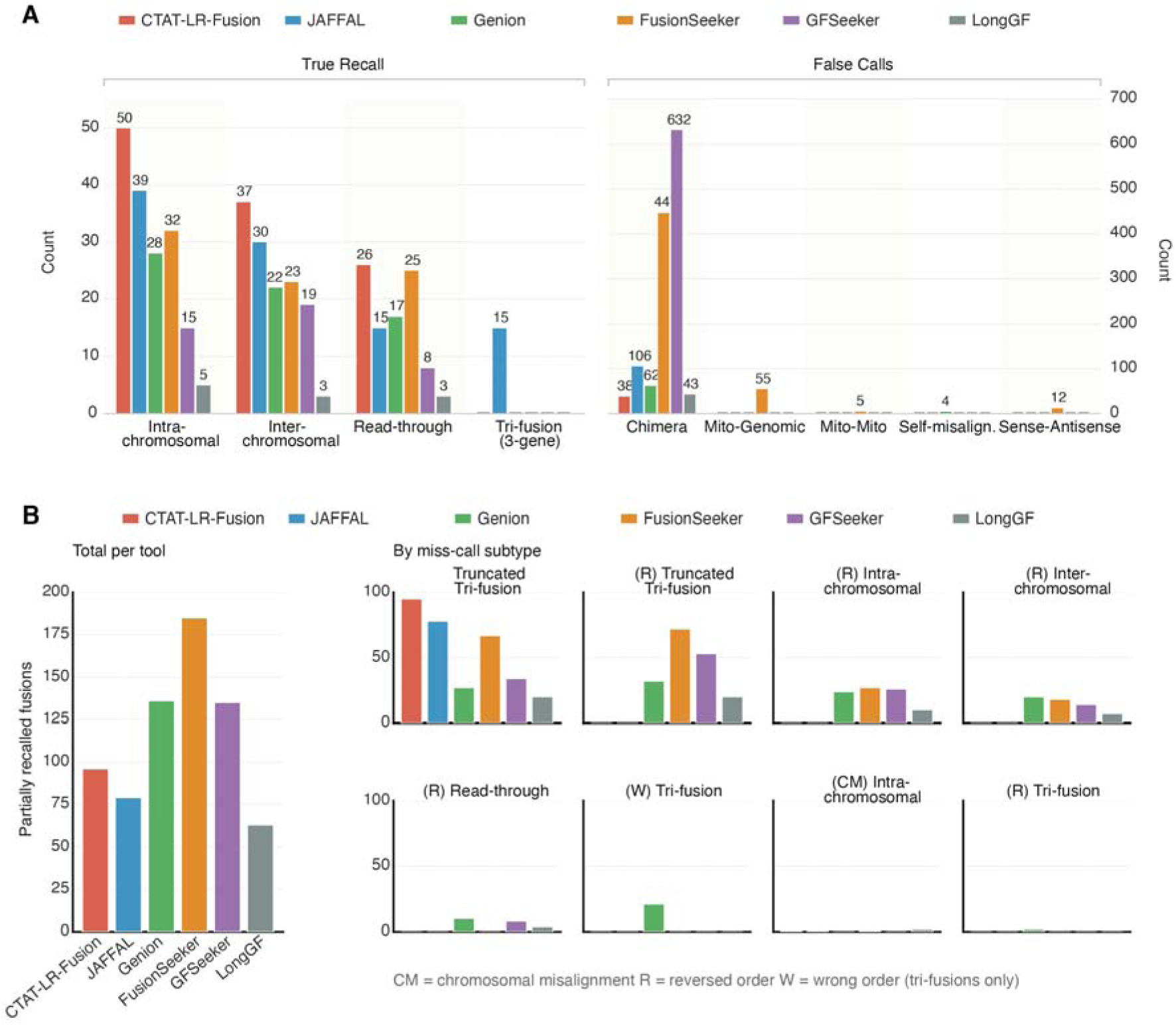
False calls and partially recalled fusions. **(A)** The bar plots show the number of recalled fusions for each tool and false fusions by type and tool. **(B)** The total number of partially called fusions for each tool and miscalls by type. M = Chromosomal-misalignment, R = Reversed order, W = Wrong Order (applicable to tri-fusions only); 100 Gb, 95% identity.

**Supplementary Figure 6.**
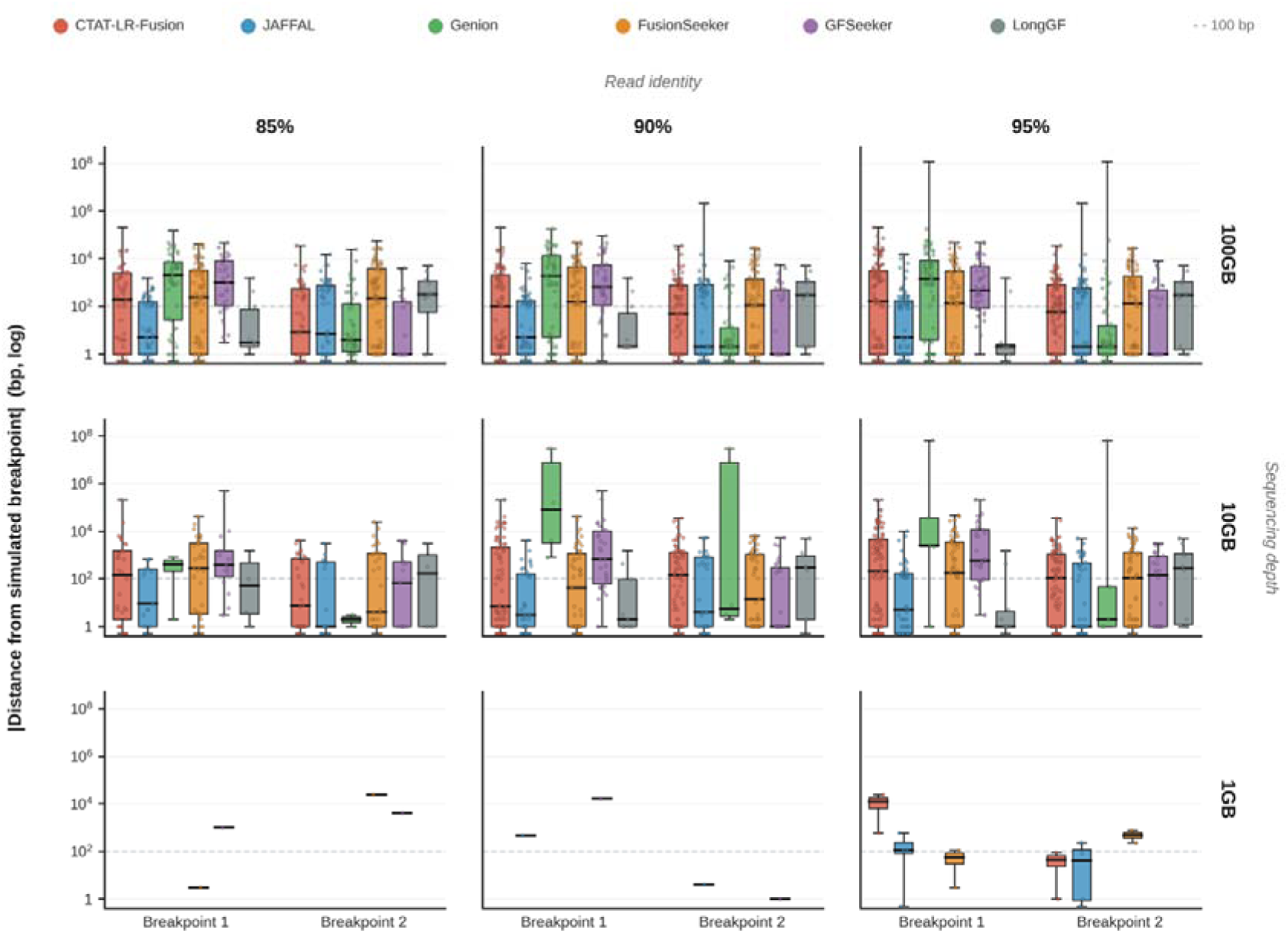
The absolute distance between the breakpoint predicted by the fusion caller and the breakpoint of the simulated fusions.

**Supplementary Figure 7.**
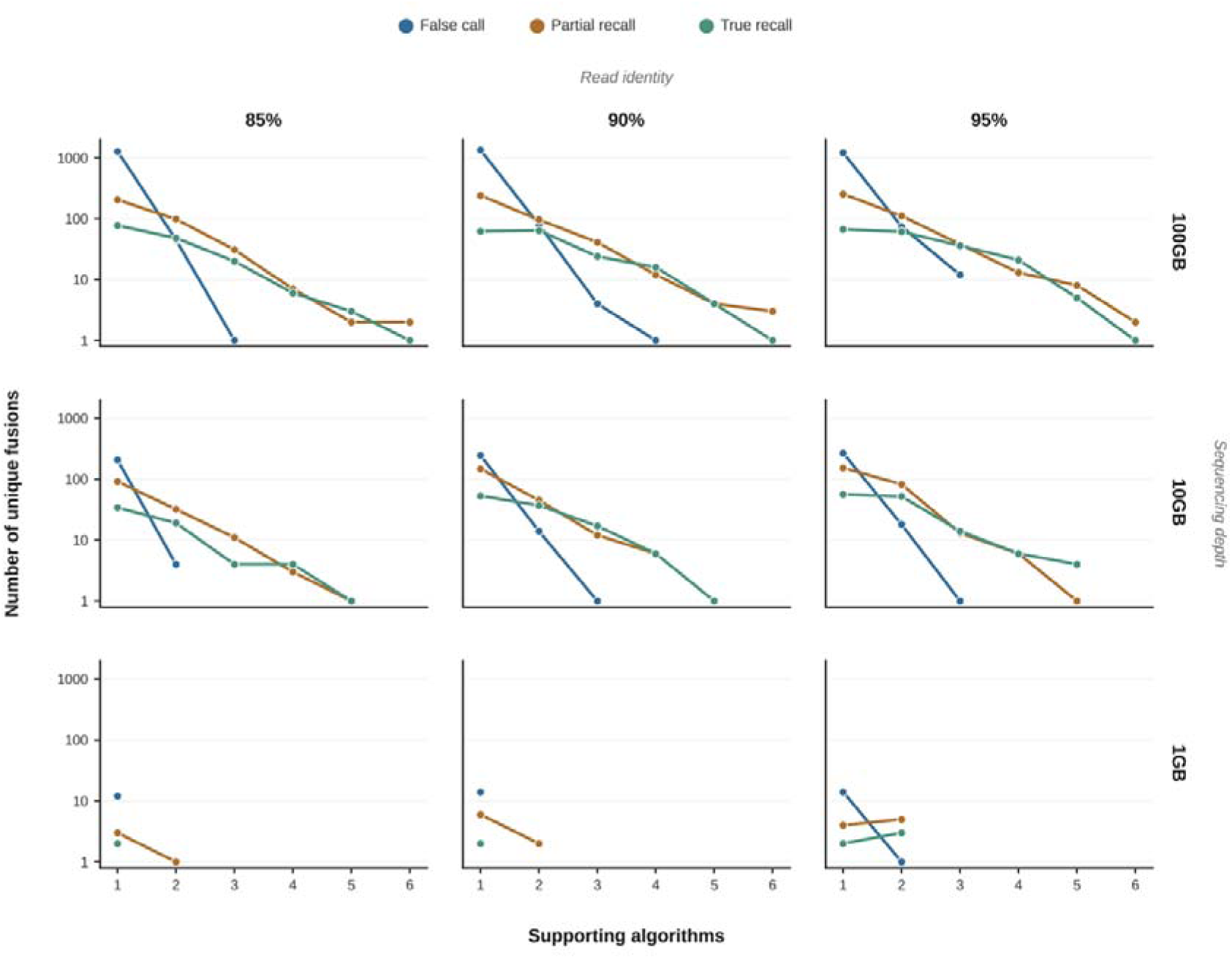
The number of unique fusions supported by 1 to 6 algorithms, 100GB 95% mean sequencing identity simulation.

**Supplementary Figure 8.**
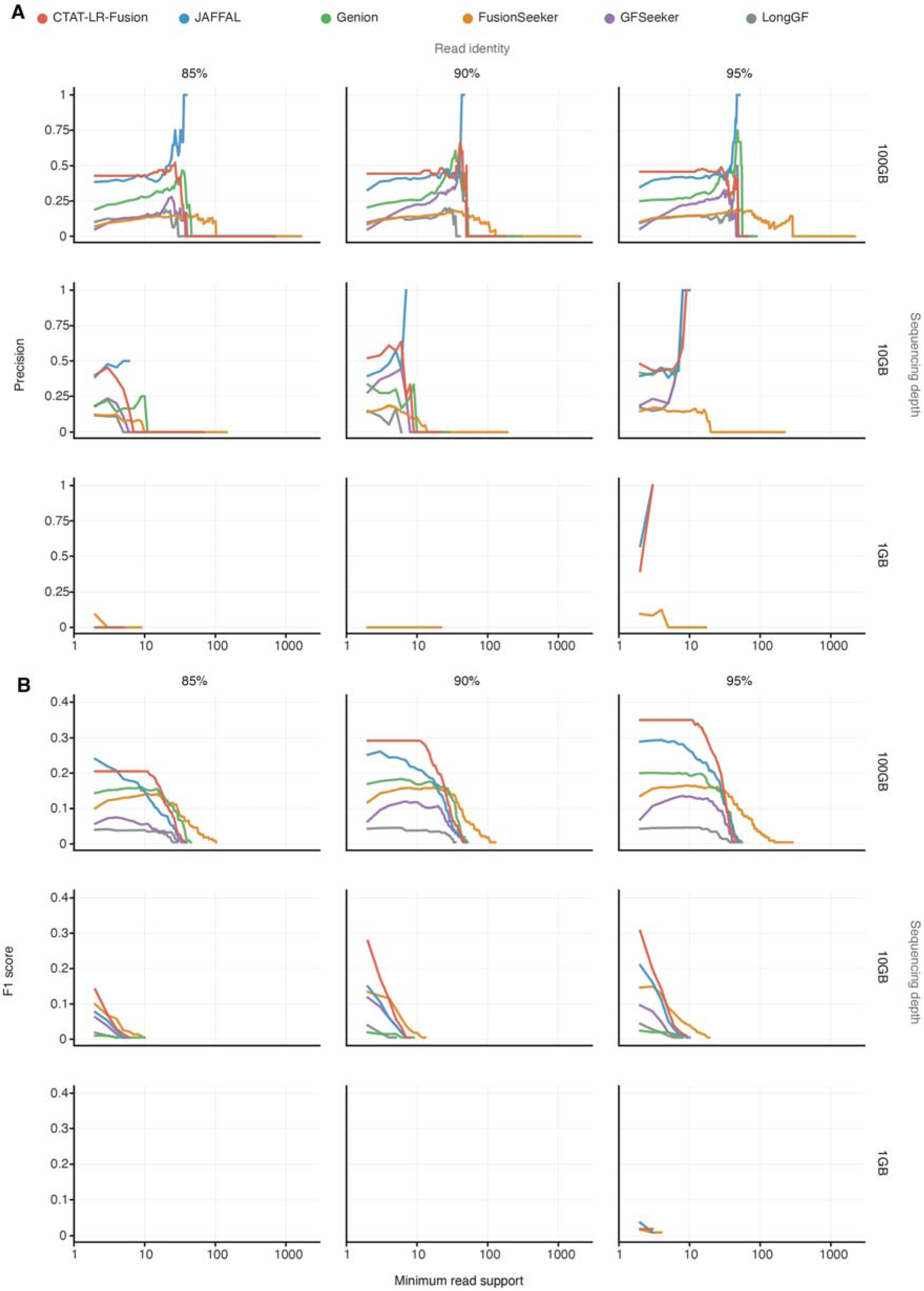
**(A)** Precision in relation to minimum read support. Partially recalled fusions were counted as false positives for calculation. **(B)** F1 score in relation to the minimum read support threshold. F1 score is a metric that combines precision by recall (2(Precision × Recall) / (Precision + Recall)). Partially recalled fusions were counted as false positives for calculation.

**Supplementary Figure 9.**
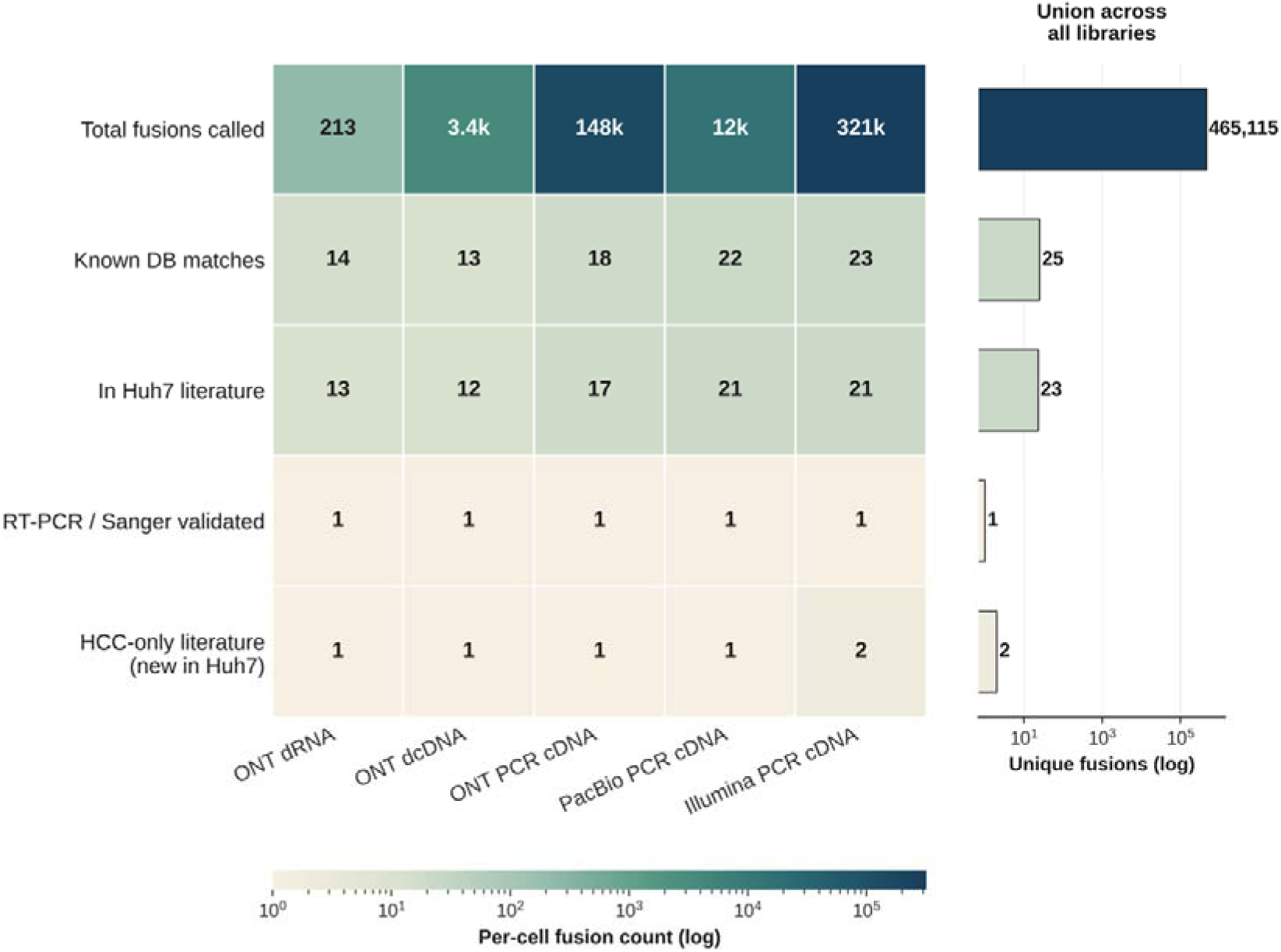
Huh7 fusion discovery summary per library and metric.

**Supplementary Figure 10.**
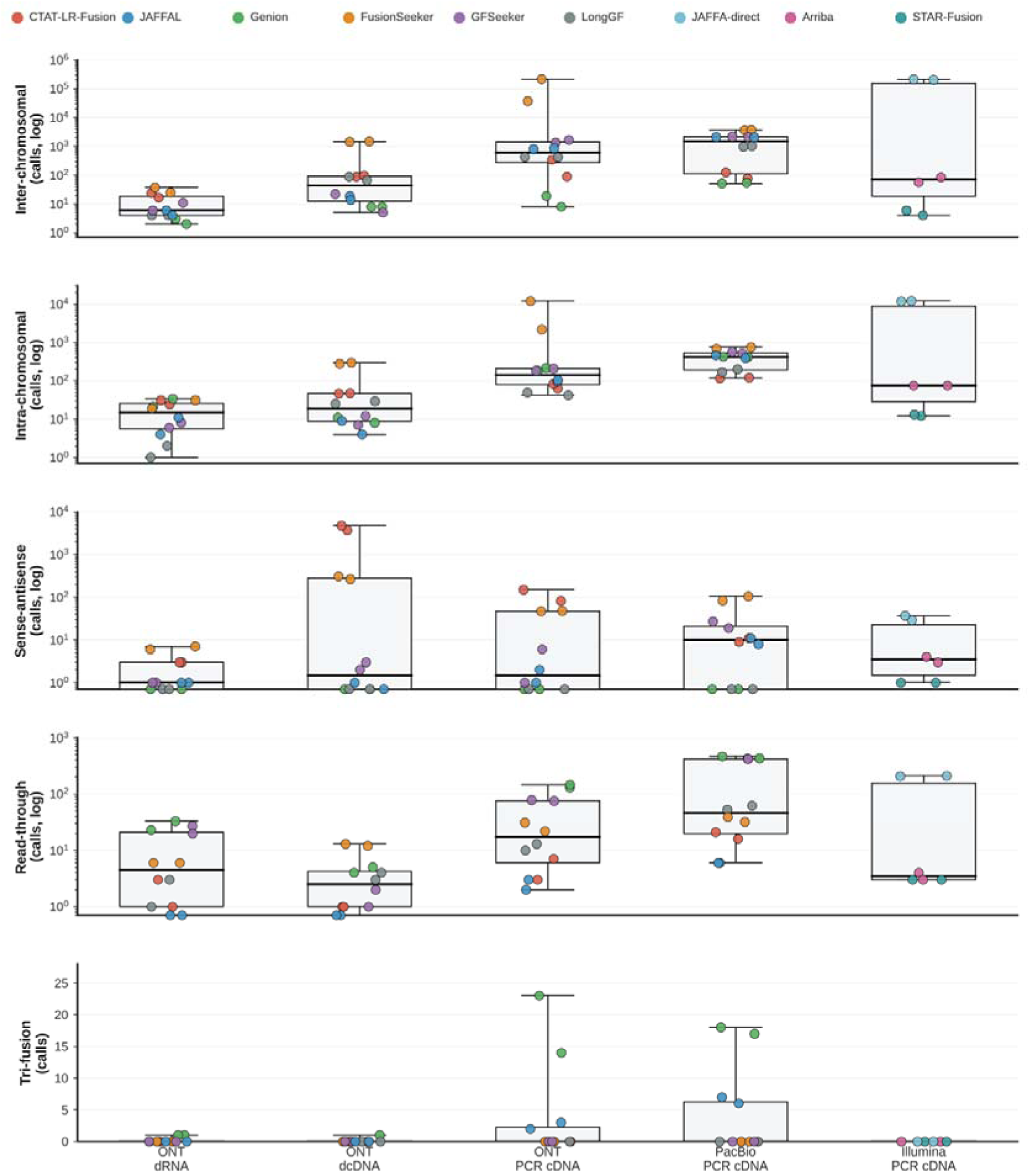
Huh7 fusion calls by fusion type across libraries.

**Supplementary Figure 11.**
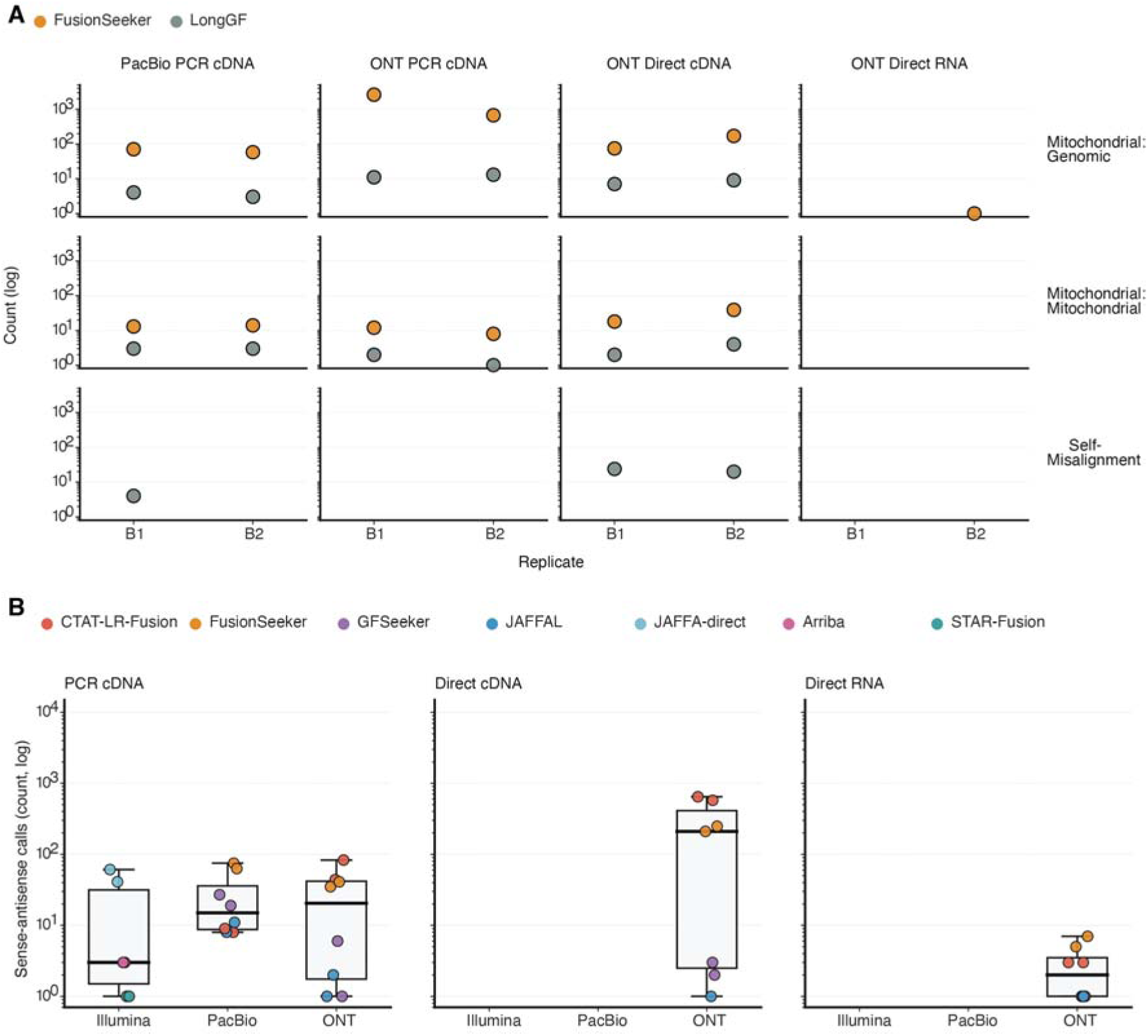
**(A)** Atypical fusion calls in Huh7 cells. Only two tools, *FusionSeeker* and *LongGF*, report mitochondrial-related chimeras. **(B)** Sense-antisense fusion calls in Huh7.

**Supplementary Figure 12.**
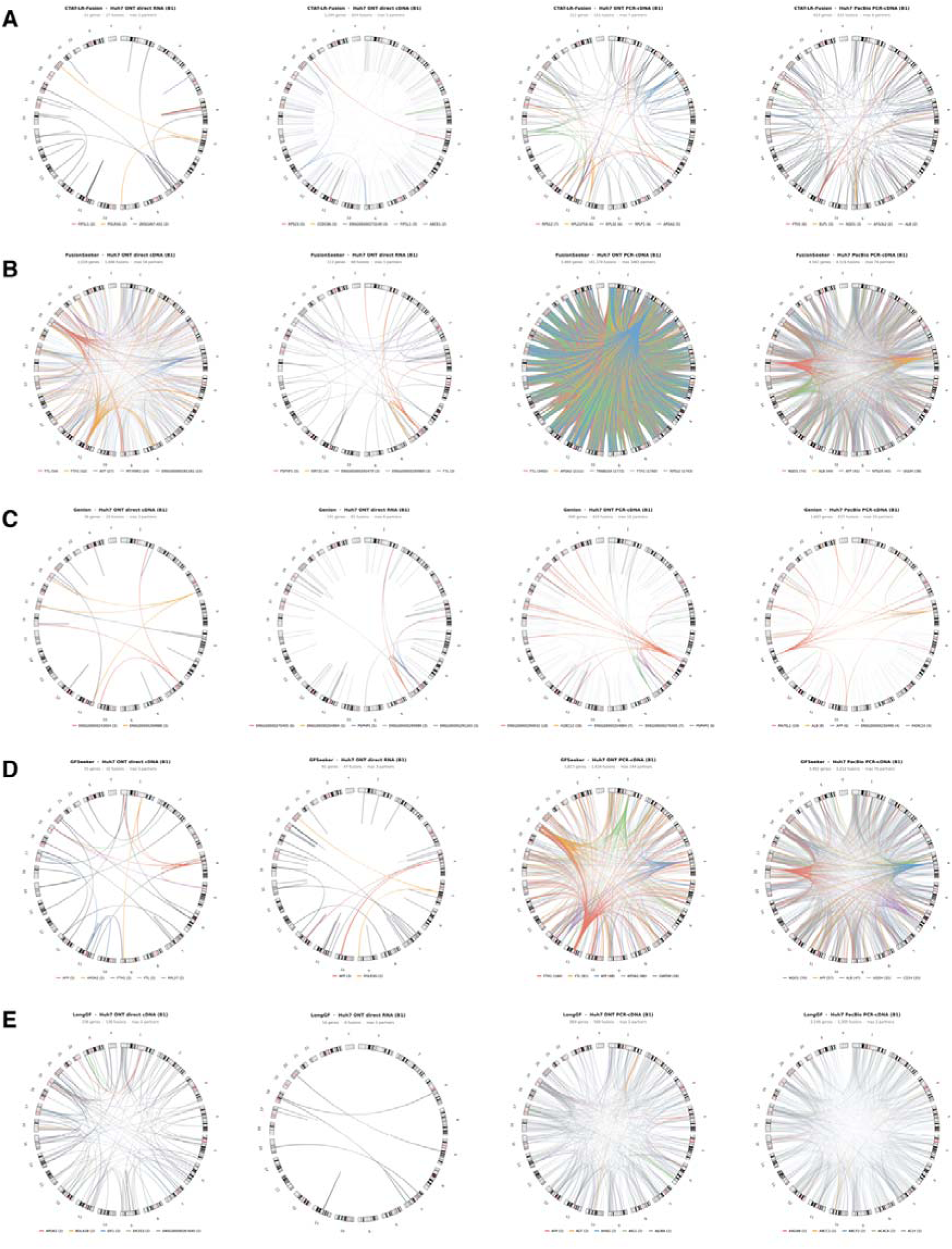
Circus plots illustrating fusion partner promiscuity. Fusion-partner networks with chromosomes drawn as hg38 cytoband ideogram; each arc represents a fusion transcript, with the top five hub genes coloured (others are grey). **(A)** CTAT-LR-Fusion; **(B)** FusionSeeker; **(C)** Genion; **(D)** GFSeeker; **(E)** LongGF.

**Supplementary Figure 13.**
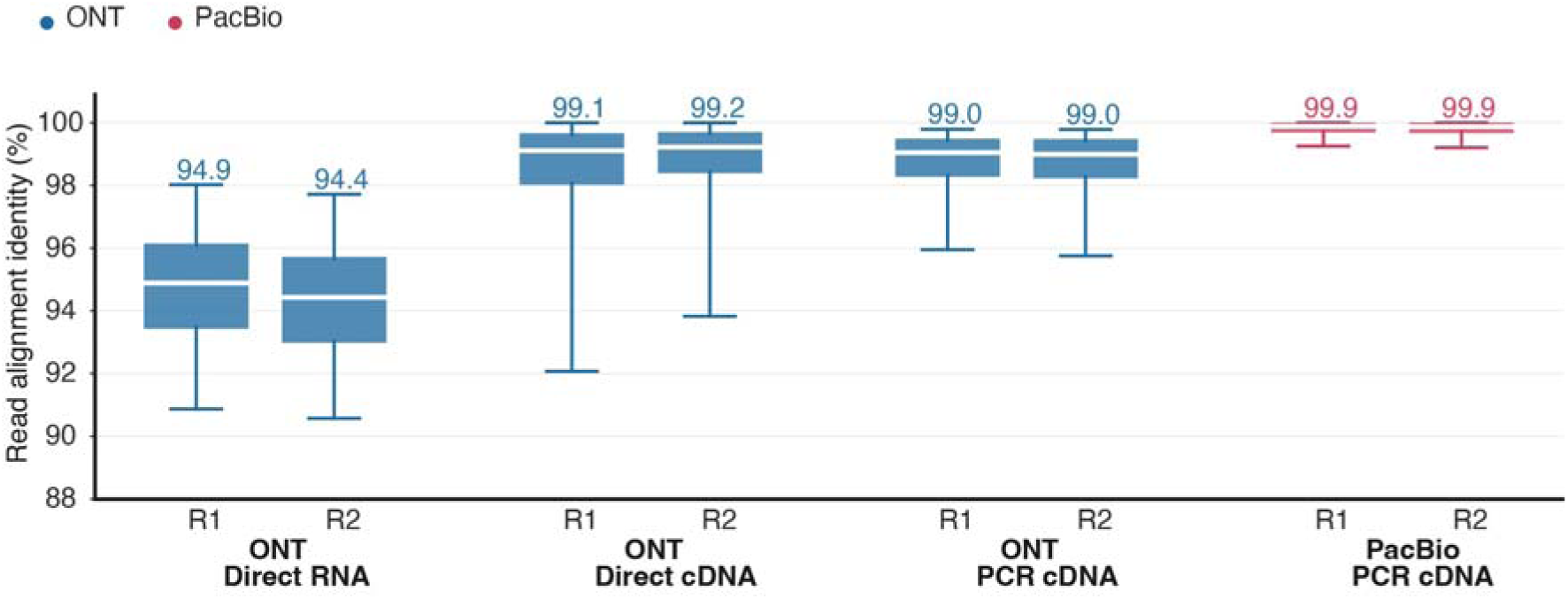
Read alignment identity by library preparation. Distributions of per-read alignment identity for the four long-read Huh7 library preparations: ONT direct RNA, ONT direct cDNA, ONT PCR-cDNA and PacBio PCR-cDNA. Alignment identity is defined as (aligned bases minus edit distance) divided by aligned bases, with introns and soft or hard clips excluded so that spliced alignments are not penalised.

**Supplementary Figure 14.**
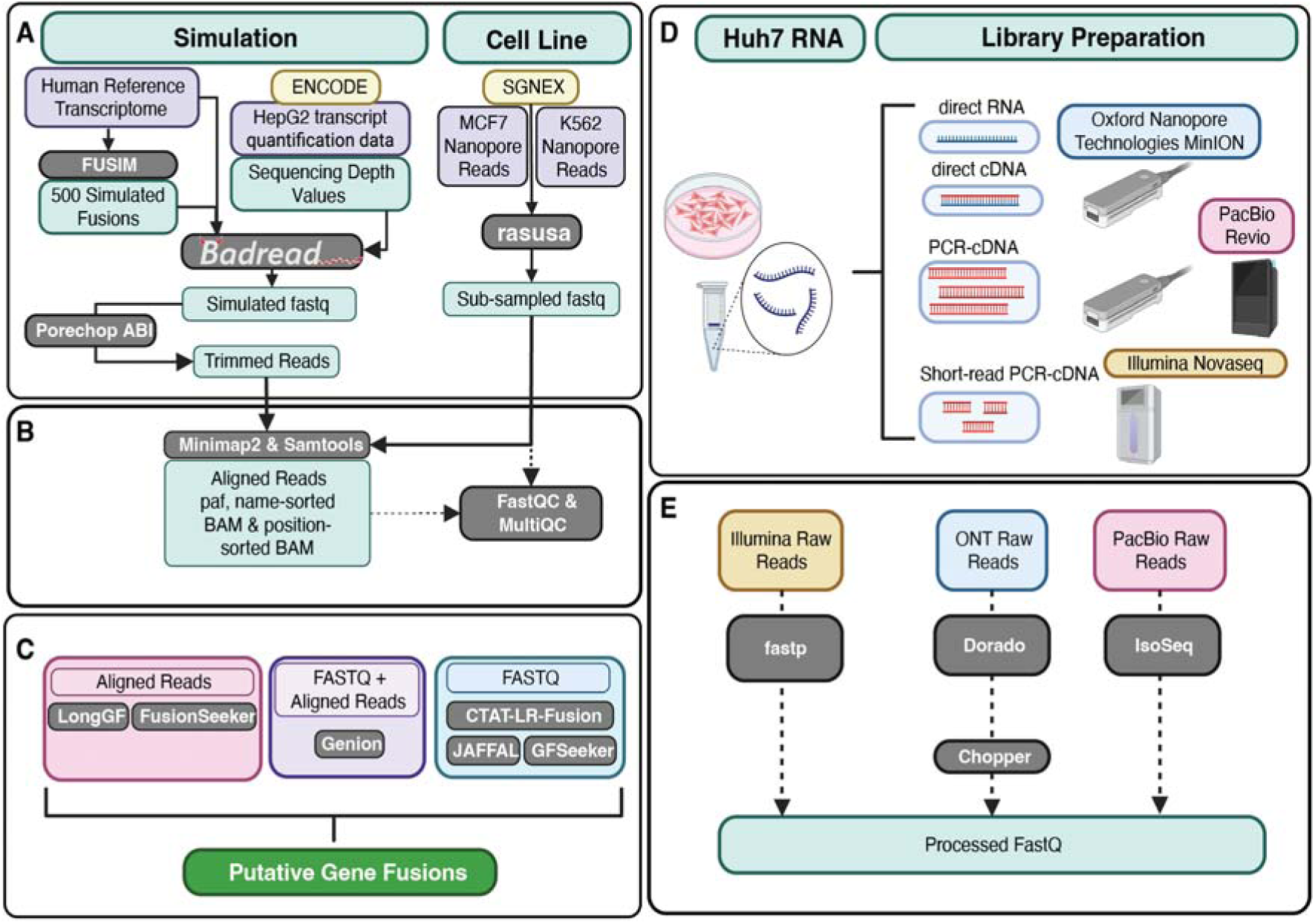
Benchmarking framework implementation. **(A)** Simulated and cell line-based fusion transcripts: Simulated reads were generated using *Badread*, based on transcript quantification data from ENCODE (HepG2) and a fusion gene set created by FUSIM combined with the human reference transcriptome. Oxford Nanopore sequencing reads from cell lines K562 & MCF7 were obtained from SG-NEX and subjected to subsampling using *rasusa*. **(B)** Read alignment and quality control: All reads underwent quality control checks using *FastQC* and *MultiQC*. Simulated reads were additionally processed with *Porechop ABI* for adapter trimming. Reads were aligned to the genome using *minimap2* and *samtools*. **(C)** Fusion calling: Aligned reads were then analysed with *Genion*, *LongGF*, *FusionSeeker*, *JAFFAL*, *CTAT-LR-Fusion*, and *GFSeeker*. **(D)** Huh7 cell transcriptomes were sequenced with multiple platforms and library preparation methods, including ONT direct RNA, direct cDNA, and PCR-cDNA sequencing, PacBio Kinnex sequencing, and Illumina short-read sequencing. **(E)** Huh7 sequencing data were preprocessed with *fastp* (for Illumina short-reads), *IsoSeq3* (for PacBio long-reads), *Dorado* (ONT long-read basecalling).

## Notes

### Competing Interest Statement

The authors have declared no competing interest.

